# Conserved and divergent features of neuronal CaMKII holoenzyme structure, function, and high-order assembly

**DOI:** 10.1101/2021.01.21.427643

**Authors:** Olivia R. Buonarati, Adam P. Miller, Steven J. Coultrap, K. Ulrich Bayer, Steve L. Reichow

**Author notes:** equal contribution authors, listed in alphabetical order (Olivia R. Buonarati, Adam P. Miller).

## Abstract

Neuronal CaMKII holoenzymes (α- and β-isoforms) enable molecular signal computation underlying learning and memory, but also mediate excitotoxic neuronal death. Here, we provide a comparative analysis of these signaling devices, using single particle EM in combination with biochemical and live-cell imaging studies. In the basal state, both isoforms assembled mainly as 12-mers (but also 14-mers, and even 16-mers for the β-isoform). CaMKIIα and β-isoforms adopted an ensemble of extended activatable states (with average radius of 12.6 versus 16.8 nm, respectively), characterized by multiple transient intra- and inter-holoenzyme interactions associated with distinct functional properties. The extended state of CaMKIIβ allowed EM analysis to directly resolve intra-holoenzyme kinase-domain dimers that could enable the cooperative activation mechanism by calmodulin, which was found for both isoforms. Surprisingly, high-order CaMKII clustering mediated by inter-holoenzyme kinase-domain dimerization was reduced for the β isoform for both basal and excitotoxicity-induced clusters, both *in vitro* and in neurons.

## INTRODUCTION

The Ca^2+^/calmodulin(CaM)-dependent protein kinase II (CaMKII) is a major mediator of long-term plasticity at excitatory glutamatergic synapses in the hippocampus that is required for learning and memory (Bayer and Schulman, 2019; Hell, 2014; Lisman et al., 2012). Beyond these physiological functions, CaMKII also mediates the glutamate excitotoxicity that kills neurons during ischemia (Coultrap et al., 2011; Deng et al., 2017; Vest et al., 2010). Both synaptic plasticity and excitotoxic cell death require the 12-meric CaMKII holoenzyme structure for at least two key regulatory functions: i) autophosphorylation at T286 (pT286)(Coultrap et al., 2014; Deng et al., 2017; Giese et al., 1998), which occurs between subunits within a holoenzyme (Hanson et al., 1994) and thereby enables detection of the stimulation frequency (De Koninck and Schulman, 1998); and ii) both long-term potentiation (LTP) and ischemic cell death require CaMKII binding to the NMDA-type glutamate receptor subunit GluN2B (Barria and Malinow, 2005; Buonarati et al., 2020; Halt et al., 2012), which also requires the holoenzyme structure (Bayer et al., 2006; Strack et al., 2000) and then mediates CaMKII accumulation at synapses during LTP and excitotoxic insults (Bayer et al., 2001; Buonarati et al., 2020; Halt et al., 2012). Both pT286 and GluN2B binding require an initial stimulus by Ca^2+^/CaM, but then maintain partial “autonomous” kinase activity even after Ca^2+^/CaM has dissociated (Bayer et al., 2001; Bayer and Schulman, 2019; Miller and Kennedy, 1986). By contrast, the Ca^2+^/CaM-induced clustering of multiple holoenzymes into higher order aggregates is thought to restrict kinase activity (Hudmon et al., 1996). This aggregation requires ischemia-related conditions (such as low pH and higher ADP than ATP concentration) and mediates the extrasynaptic clustering in response to excitotoxic stimuli (Dosemeci et al., 2000; Hudmon et al., 1996; Hudmon et al., 2001; Vest et al., 2009), but may contribute also to the synaptic accumulation in response to LTP stimuli (Hudmon et al., 2005).

Together, these holoenzyme functions are thought to provide essential mechanisms for information processing and storage (Bayer and Schulman, 2019; Coultrap and Bayer, 2012b; Rossetti et al., 2017). Thus, elucidating the CaMKII holoenzyme structure that enables these mechanisms has been of long-standing interest, with the first electron microscopy (EM) studies performed over 30 years ago (Kanaseki et al., 1991; Woodgett et al., 1983). CaMKII holoenzymes are oligomeric assemblies, with each subunit containing an N-terminal kinase domain, followed by a variable internal linker region that connects to a C-terminal association (or hub) domain that is responsible for oligomerization. More recently, a high-resolution crystal structure showed the 12-meric holoenzyme in a compact conformation, with the N-terminal kinase domains folding back onto the association domain that forms a central hub complex (Chao et al., 2011). Notably, this compact conformation is not activation-competent (as the CaM binding regulatory region is buried), and transition between the compact and an extended conformation provided a potential regulatory mechanism for cooperative activation by CaM. However, subsequent studies indicated that the vast majority of kinase domains is in the activation-competent extended conformation (>95%), both *in vitro* and in intact cells (Myers et al., 2017; Sloutsky et al., 2020), indicating that cooperativity must be mediated by a different mechanism.

To gain deeper insight into the conserved structural and functional features of CaMKII holoenzymes, we performed a comparative a single-particle EM analysis to CaMKIIβ, the second most prevalent isoform in neurons (Bayer et al., 1999; Bennett and Kennedy, 1987; Cook et al., 2018; Tombes et al., 2003). As we have previously described for the α isoform (Myers et al., 2017), the association domains of the β isoform form a rigid central hub capable of adopting multiple stoichiometries, whereas their kinase domains where primarily extended away from the hub in an activation-competent and highly flexible manner (see Figure 1). Within the ensemble of conformational states detected by single particle analysis, our study revealed several intriguing similarities and differences between these isoforms, including formation of higher order holoenzyme clusters, which are thought to form via dimerization of kinase domains from different holoenzymes and increase in response to ischemic or excitotoxic insults (Hudmon et al., 2001; Vest et al., 2009). In addition, we found the first direct evidence for dimer formation between kinase domains of the same holoenzyme, a structural feature that can mediate the cooperative activation by CaM found here for both CaMKIIα and β isoforms. Thus, our analysis of CaMKIIβ revealed not only isoform-specific differences, but also generally conserved themes of CaMKII regulation that are facilitated by a dynamic ensemble of multiple transient interactions supported by the holoenzyme architecture.

**Figure 1.**
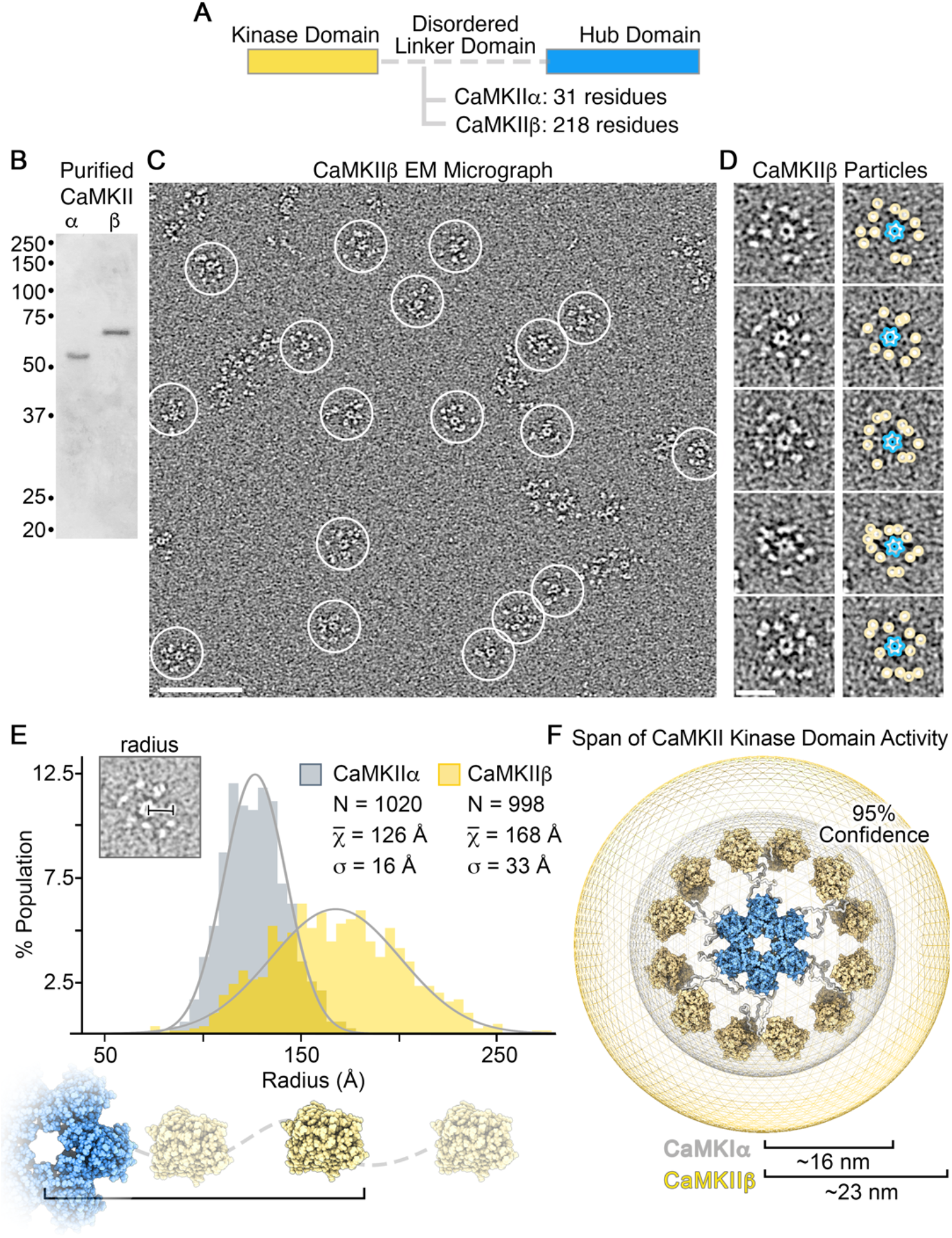
Comparative structural analysis of CaMKII holoenzymes resolved by single particle EM. (A) Diagram showing the major difference in CaMKIIα and CaMKIIβ domain architecture is within the length of their respective disordered linker domains. (B) SDS-PAGE gel showing full-length CaMKIIα and CaMKIIβ isoforms purified from S*f*9 cells migrate at the expected molecular weights and display no sign of proteolytic degradation. (C) Electron micrograph of negatively stained CaMKIIβ holoenzymes. Individual complexes are indicated by white circles. Scale bar = 100 nm. (D) Zoom-view of individual particles with the hub complex (blue outline) and resolved kinase domains (yellow circle) indicated. Scale bar = 20 nm. (E) Histogram analysis of kinase radius (center of hub to outer radius of kinase domain) obtained for CaMKIIα (grey, n=1020) and CaMKIIβ (yellow, n=998) particles, bin = 5 Å. Grey lines represent a gaussian fit to the data. *Inset*, illustrating radii were obtained from raw particle images. (F) Schematic illustrating the difference in radial expansion sampled by CaMKIIα (grey) versus CaMKIIβ (yellow), where the outer boundary and values (16 nm and 23 nm, respectively) represents the 95% confidence interval of measured holoenzyme radii.

## RESULTS

### The dodecameric CaMKIIβ holoenzyme adopts an extended kinase radius

We have shown previously that the CaMKIIα holoenzyme adopts a predominant dodecameric (12-mer) assembly, with an extended and flexible activatable-state conformation as visualized by negative stain electron microscopy (NSEM) (Myers et al., 2017). This 12-meric assembly is organized by a central and well-ordered hub domain complex, with 12 kinase domains displayed in an extended fashion via an intrinsically disordered flexible linker domain (Figure 1A). For comparative structural analysis to the β isoform, CaMKIIβ holoenzymes were expressed in eukaryotic cells (S*f*9), purified by chromatographic methods and prepared for NSEM using the same protocols previously described for CaMKIIα. Isolated holoenzymes show no signs of proteolytic degradation by SDS-PAGE (Figure 1B), and CaMKIIβ specimens produced well-resolved assemblies resembling the same “flower-like” appearance of CaMKIIα holoenzyme structures observed by NSEM (Figure 1C,D). A defined central ring of density corresponding to the hub domain (~11 nm diameter) was clearly resolved, surrounded by an array of smaller densities, “the petals,” corresponding to tethered kinase domains (diameter ~5 nm) (Figure 1C,D). However, despite an overall structural similarity, the peripheral kinase domains associated with CaMKIIβ holoenzymes were qualitatively more extended and heterogeneously configured around the hub domain, as compared to CaMKIIα holoenzymes. Although the disordered linker domain is not resolvable by NSEM, these differences are consistent with the primary divergence in amino acid sequence between CaMKII isoforms, where in CaMKIIβ the linker domain is ~218 residues long (as compared to ~31 residue linker in CaMKIIα) (Cook et al., 2018; Tombes et al., 2003) (Figure 1A).

For both CaMKIIα and CaMKIIβ, the flexible linker region facilitates the formation of a variety (or continuum) of conformational states that are not directly amenable to traditional EM image classification and averaging methods. Therefore, for quantitative characterization and comparative analysis between holoenzyme structures, we took advantage of the high-contrast of NSEM imaging to conduct a series of measurements and statistical analyses conducted directly on individual holoenzyme particle images obtained from raw micrographs (Myers et al., 2017) (Figure 1D,E). In the first set of measurements, the radial extension for each kinase domain (*i.e*., kinase radius) was obtained by measuring from the center of the hub complex to the center of each kinase domain, and this measurement was then appended by 2.25 nm to account for the approximate radius of the kinase domain (Figure 1E,F). For both isoforms, the distribution of kinase domain radii obtained from ~1000 particle measurements for each isoform appears randomly positioned with apparent gaussian distribution (Figure 1E). However, the average kinase radius of CaMKIIβ is significantly larger at ~16.8 nm (± 0.1 SEM) as compared to CaMKIIα assemblies ~12.6 nm (± 0.05 SEM) (P < 0.001). Although the linker domain is not directly visible by NSEM, the edge-to-edge distance separating the kinase domain and hub domain may be used as an approximation of the linker extension. These calculations provide an average linker extension of ~2.7 nm for CaMKIIα and ~6.8 nm for CaMKIIβ. These values are consistent with random chain polymer models (traditional random walk model) based on the differences in amino acid lengths of the respective linker domains (Flory, 1975), and further support the idea that CaMKII kinase domains are freely tethered to the central hub domain by intrinsically disordered linker regions. Based on these measurements, a 95% confidence of kinase domain positioning can be expected to span a radius of up to ~16 nm for CaMKIIα holoenzymes and up to ~23 nm for CaMKIIβ holoenzymes (Figure 1F).

CaMKII holoenzymes can adopt a ‘compact state’ involving kinase-hub domain interaction, but only a minor fraction of CaMKIIα holoenzymes was found in this conformation (Chao et al., 2011; Myers et al., 2017; Sloutsky et al., 2020). From our analysis in Figure 1E, a kinase domain radius measured less than ~10 nm would potentially place a kinase domain in steric contact with the central hub complex. Consistent with our previous analysis, CaMKIIα holoenzymes showed only a small fraction of individual subunits with kinase domain radii that fall within this category (~3% of kinase domains with radius < 10 nm) ((Myers et al., 2017) and Figure 1E). In comparison, CaMKIIβ displayed less than 1% of kinase domains with radius < 10 nm observed by NSEM (Figure 1E), indicating that a ‘compact state’ is at most only sparsely populated.

### Autophosphorylation of pT286 in CaMKIIα versus pT287 in CaMKIIβ

The extended radius of the CaMKIIβ compared to the α holoenzyme would lead to a lower local concentration of kinase domains within the space occupied by a holoenzyme, based on simple geometric considerations (~1.0 mM versus ~2.3 mM; see Figure S1). Thus, we decided to directly compare CaMKIIα versus β purified holoenzymes for autophosphorylation at T286 (in α) or T287 (in β), which occurs as an inter-subunit intra-holoenzyme reaction (Bradshaw et al., 2002; Hanson et al., 1994; Rich and Schulman, 1998). *In vitro* kinase stimulation with Ca^2+^/CaM resulted fast autophosphorylation for both CaMKIIα and β, with no significant differences between T286 and T287 detected throughout the reaction time-course (Figure S1). Taken together, these results indicate that the significant difference in the proximity of kinase domains in CaMKIIβ versus CaMKIIα does not significantly affect autophosphorylation kinetics of the holoenzymes.

### CaMKIIβ forms multimeric assemblies of 12 – 16-mers

Early EM studies had already indicated that CaMKIIα forms mainly 12-mers, but some studies suggested a smaller number of subunits, particularly for the CaMKIIβ isoform (Kanaseki et al., 1991; Woodgett et al., 1983). Thus, we decided to further compare the stoichiometry of CaMKIIα versus β holoenzymes, specifically by quantifying the symmetry of their hub domain assemblies. We performed focused 2D image classification on the NSEM datasets for both isoforms by applying a circular mask to remove signal from the peripheral kinase domains and focus the image alignment procedures on the central hub domain (Figure 2A,B). The results of this analysis showed a predominance of particles displaying 6-fold radial symmetry, with dimensions and structural features matching 2D-projections the dodecameric (12-mer) hub assembly, representing 88% of the population for CaMKIIα and ~92.7% for CaMKIβ (Figure 2A–D). For both isoforms, a smaller but significant population of hub domains displaying 7-fold radial symmetry were also observed, corresponding to ~5.2% of the population for CaMKIIα and ~5.5% for CaMKIIβ holoenzymes (Figure 2A–D). The dimensions of 7-fold symmetric hubs class averages were also consistent with 2D-projections of a previously published crystallographic model of an isolated tetradecameric (14-mer) hub assembly (Hoelz et al., 2003) (Figure 2A–C).

**Figure 2.**
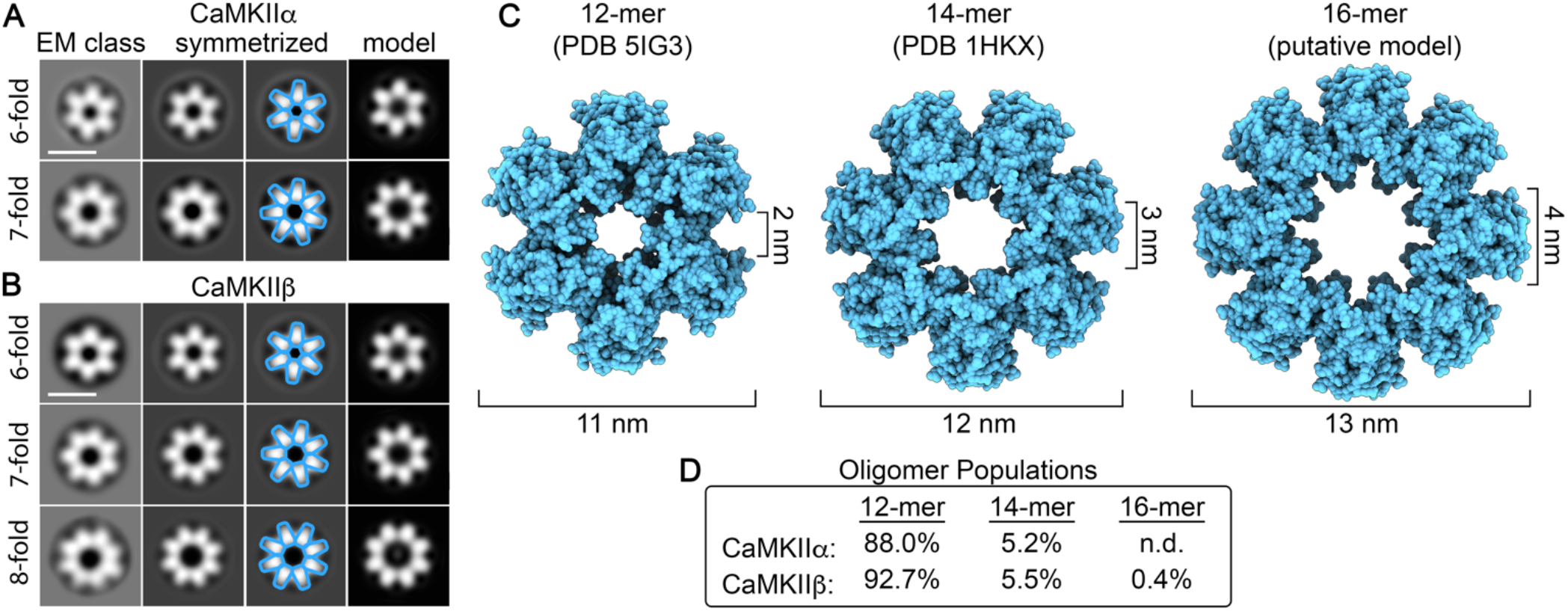
Comparative analysis of CaMKII holoenzyme stoichiometries resolved by single particle EM. (A and B) Single particle EM image classification and analysis of the central hub domain of CaMKIIα and CaMKIIβ holoenzymes, respectively. *Left*, Focused EM class averages obtained using an applied image mask (15 nm outer diameter) and without applied symmetry. *Middle left and right*, display symmetrized versions of the EM classes as indicated and with resolved features annotated (blue outline). *Right*, displays 2D back-projections from atomic models of hub assemblies displayed in panel (C), filtered to 30 Å. Scale bar = 10 nm. (C) *Left*, atomic model of a dodecameric (12-mer) hub domain (blue surface representation, PDBID 5IG3 (Bhattacharyya et al., 2016), *center*, heptadecameric (14-mer) hub domain (PDBID 1HKX (Hoelz et al., 2003), and *right*, pseudo-atomic model of the putative hexadecameric (16-mer) hub domain resolved in panel B. (D) Population of CaMKII holoenzyme stoichiometries resolved by single particle EM. For the CaMKIIα dataset n=10,902 and for the CaMKIIβ dataset n=17,347. The 16-mer was not detected (n.d.) in the smaller CaMKIIα image dataset.

Remarkably, an additional minor population of hub domain structures was detected with clear 8-fold symmetry in the CaMKIIβ image dataset, constituting ~0.4% of the population (Figure 2B–D). The dimensions of the detected 16-mer hub are ~13 nm in outer diameter and central pore measuring ~4 nm in diameter (Figure 2B). Pseudo-atomic modeling of hub domain subunits, restricted by the dimensions of the 8-fold symmetric projection average, show a reasonable fit, with minimal steric overlap between neighboring subunits, resulting in a putative hexadecameric (16-mer) hub model (Figure 2C). This pseudo-atomic model produces 2D back-projections matching the experimental density (Figure 2B). 16-mer assemblies were not detected in the CaMKIIα dataset. However, given the small population of 16-mers observed for CaMKIIβ, it is possible that this species was simply not detectable by image classification due to the significantly smaller image dataset obtained for CaMKIIα. Nevertheless, inherent differences between isoforms cannot be ruled out.

It is also noteworthy that smaller assemblies (*e.g*., 8 – 10mers) were not detected in either dataset. Given the ability of our focused image classification approach to detect populations that represent < 1% of species present, it is likely that the CaMKII hub domain is not capable of supporting such configurations under basal-state conditions (at least to any appreciable degree).

### Resolution of kinase-domain dimers within intact CaMKIIβ holoenzymes

Kinase-kinase domain pairing interactions have been proposed as a potential mechanism for CaMKII cooperative activation by Ca^2+^/CaM. Indeed, such dimers have been observed in crystals of the kinase domain (Rosenberg et al., 2005), however, they have not yet been directly detected in context of the intact holoenzyme. To assess for the presence of kinase domain dimerization in the context of CaMKII holoenzymes, the separation distance (center-to-center) between nearest neighbor kinase domains were measured on individual particle EM images (Figure 3A). For CaMKIIα holoenzymes, the average separation distance was 5.9 nm (± 0.05 SEM), with a near gaussian distribution (Figure 3A, grey). This value is consistent with our previous analysis and with solution-state FRET studies conducted on CaMKIIα holoenzymes (Myers et al., 2017; Thaler et al., 2009). These measurements indicate that the majority of kinase domains are non-interacting (Myers et al., 2017). However, an appreciable fraction of kinase separation distances (~19%) measured less than 4.5 nm (*i.e*., within steric contact distance), indicating a potential for kinase domain pairing within the context of the holoenzyme. For CaMKIIβ holoenzymes, the average kinase separation distance is significantly larger 9.4 nm (± 0.2 SEM) (Figure 3A, yellow), presumably facilitated by the longer linker domain present in this isoform. Notably, the distribution is skewed from a random gaussian distribution, toward shorter separation distances, with the most populated distance bin of 4.5 – 5.0 nm (Figure 3A). The observed deviation from random distribution toward shorter separation distances may reflect intrinsic interactions between neighboring kinase domains. In fact, while these data are consistent with the majority of kinase domains adopting a non-interacting configuration, a significant fraction of CaMKIIβ kinase domains (~8%) are also localized within our designated threshold for steric contact distance of less than 4.5 nm, indicating that kinase-domain pairing may represent a significant population of both CaMKIIα and CaMKIIβ holoenzyme structures.

**Figure 3.**
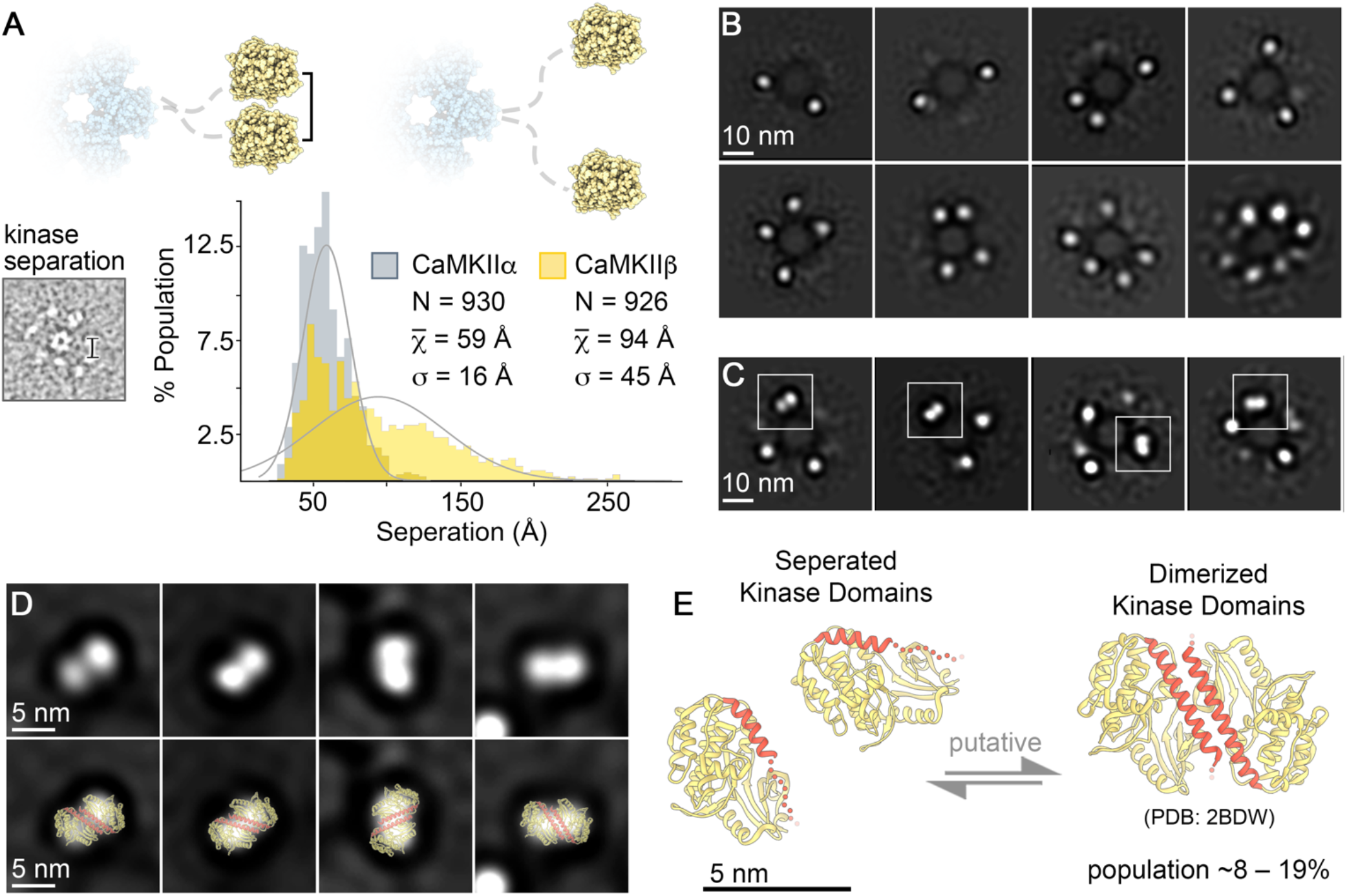
Kinase domain dimers resolved in the CaMKIIβ by single particle EM. (A) Histogram of measured distance separating neighboring kinase domains (center-to-center) obtained for CaMKIIα (grey, n=930) and CaMKIIβ (yellow, n=926) particles, bin = 5 Å. Grey lines represent a gaussian fit to the data. *Inset*, illustrating distances were obtained from raw particle images in EM micrographs. (B and C) Single particle EM image classification and analysis of CaMKIIβ holoenzymes with an applied mask to remove contribution of the hub domain during the alignment procedure (13.5 nm diameter). Panel (B), isolated kinase domains appear as punctate densities of ~4–5 nm in diameter. Only a subset of all 12 kinase domains could be resolved in class averages (typically 2 to ~7), due to spatial heterogeneity of kinase domain positions present within the population of CaMKIIβ holoenzymes. Panel (C), class averages displaying additionally resolve bilobed densities with approximate dimensions of ~10 x 5 nm (indicated by white squares). Scale bar = 10 nm in panels B and C. (D) *Top row*, zoom view of bilobed densities resolved in class averages in panel (C), and *bottom row*, with fitted crystallographic structure of the *C. elegans* CaMKIIα kinase/regulatory domain (yellow/red ribbon; PDBID 2BDW (Rosenberg et al., 2005) previously shown to form a dimeric interface involving the regulatory domain (red). (E) Putative model depicting an equilibrium of states involving independent kinase domains (PDBID 2VZ6) (Rellos et al., 2010) and kinase domain dimer (PDBID 2BDW) (Rosenberg et al., 2005) proposed to be present in the context of the CaMKII holoenzyme. The regulatory domain (red) in the dimerized state becomes more ordered and occluded from CaM binding. Scale bar = 5 nm in panels D and E.

To further support this evaluation, we conducted focused 2D image classification on CaMKIIβ kinase domains, this time by masking away the central density of the hub domain prior to image alignment (Figure 3B,C). The results of this analysis produced two distinct groups of 2D class averages. The first group appears to resolve isolated densities of ~4–5 nm diameter, corresponding to a subset of the individual kinase domains belonging to a single holoenzyme (Figure 3B). Notably, all 12 kinase domains were not completely resolved in any of these 2D class averages obtained from CaMKIIβ holoenzymes, due to the continuum of kinase domain configurations supported by the extended linker region. In the second group of 2D class averages displayed in Figure 3C, larger elongated densities of ~10 x 5 nm are resolved in addition to the ~4–5 nm individual kinase domain densities. These larger densities have a distinct bi-lobed appearance and dimensions consistent with that of the crystalized kinase domain dimer structure (Figure 3D,E) (Rosenberg et al., 2005). For CaMKIIα, apparent kinase domain dimers were previously resolved in individual particle images (Myers et al., 2017); however, such structures could not be resolved by 2D classification procedures. We attribute this to limitations associated with local crowding of neighboring kinase domains present in this isoform which could interfere with image classification. Taken together, these data support the notion that CaMKII kinase domains are capable of forming dimers within the context of the holoenzyme assembly in both CaMKIIα and CaMKIIβ isoforms, and may represent ~8–19% of kinase domains organized by the CaMKII holoenzyme structure (Figure 3E).

### CaMKIIα and β differ in CaM activation constant but not in activation cooperativity

The kinase domain dimers that were found previously in a crystal of the kinase domain (Rosenberg et al., 2005) and potentially here in context of the holoenzyme (see Figure 3) are formed by low-affinity coiled-coil interactions between two regulatory domains; then binding of Ca^2+^/CaM to one regulatory domain would disrupt the interaction and thereby facilitate binding also to the other regulatory domain. This could explain the cooperative activation of CaMKII by CaM (as determined by a Hill slope of the activation curve that is greater than 1; Chao et al., 2011; Myers et al., 2017). Thus, we decided to directly compare CaM activation of CaMKIIα versus β purified holoenzymes; for this, we used our established kinase activity assay that measures phosphorylation of the peptide substrate syntide-2 (Coultrap and Bayer, 2011, 2014; Coultrap et al., 2010). Consistent with previous reports (Brocke et al., 1999; De Koninck and Schulman, 1998), CaMKIIβ was more sensitive to Ca^2+^/CaM-stimulation than CaMKIIα (here with an EC_50_ of 15 nM compared to 30 nM; Figure 4 and Figure S2). However, the Hill slope was indistinguishable between the isoforms and was determined to be ~1.6 for both (Figure 4 and Figure S2). Such Hill slopes between 1 and 2 are consistent with dimer formation of some but not all kinase domains within a holoenzyme, and would indicate that dimer formation is equal between α and β. Indeed, even though the dimer structure was resolved in 2D average classes only for CaMKIIβ but not α (see Figure 3 and Myers et al., 2017), the percentage of kinase domains that are close enough for potential dimer formation are comparable for both α and β isoforms (19% and 8%, respectively), and arguably within the degree of uncertainty based on the limitations of our approach (see Discussion).

**Figure 4.**
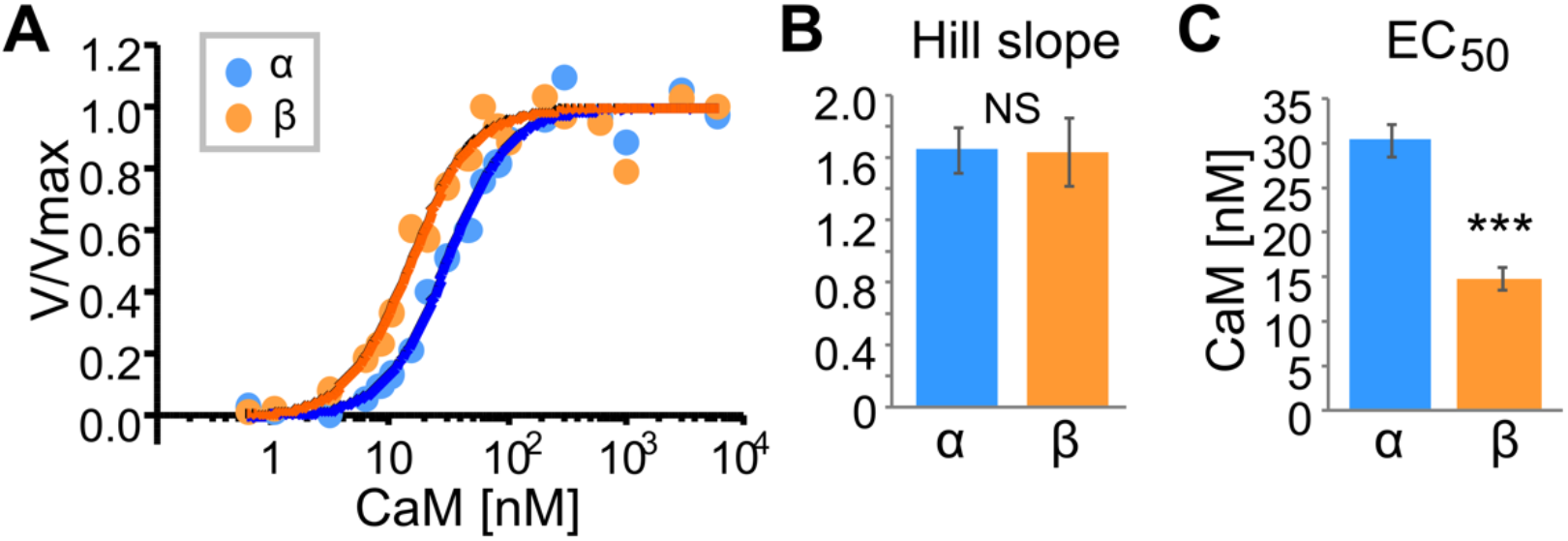
CaMKIIα and CaMKIIβ activation by CaM: different sensitivity but equal Hill coefficient. Quantifications show mean ± SEM. ***p<0.001. A) *In vitro* CaMKII activity in response to varying Ca^2+^/CaM (0.6 nM to 6 μM CaM). The curve fits shown are based on data from two independent experiments (see Fig. S2). B) Both CaMKIIα and CaMKIIβ exhibited a Hill slope coefficient greater than 1. No differences were observed between isoforms, demonstrating equal CaM cooperativity (1.65±0.15 for a and 1.64±0.22 for β; extra-sum-of-squares F-test, p=0.9668). C) As expected, CaMKIIβ showed higher sensitivity to CaM activation, with an EC_50_ significantly lower than that of CaMKIIα (14.6 and 30.1 nM, respectively; extra-sum-of-squares F-test, ***p<0.001).

### CaMKIIα and β form higher order holoenzyme clusters both *in vitro* and in neurons

Whereas the kinase domain-dimers that form within a holoenzyme may contribute to cooperative CaMKII activation, an inter-holoenzyme kinase domain-dimer formation is thought to mediate higher order clustering of CaMKII holoenzymes (although the proposed dimerization mechanisms differ; Hudmon et al., 2001; Vest et al., 2009). Some clustering can occur basally in neurons and small clusters were also observed on our EM grids, with ~56–58% of holoenzymes potentially interacting to form holoenzyme pairs and/or higher-order clusters (Figure S3). However, clustering at extrasynaptic sites is majorly enhanced by ischemic conditions (Dosemeci et al., 2000; Hudmon et al., 2005; Vest et al., 2009). Additionally, clustering may contribute to the CaMKII accumulation at excitatory spine synapses during both LTP and ischemia (Hudmon et al., 2005), although CaMKII binding to GluN2B is at least a co-requirement for both (Bayer et al., 2001; Buonarati et al., 2020; Halt et al., 2012). CaMKIIβ has been described to be incompetent for the ischemia-related clustering (at least *in vitro;* Hudmon et al., 2001), but seemed to form at least some basal clusters (see Figure S3). Thus, we decided to directly compare the clustering of CaMKIIα versus β in dissociated hippocampal neurons. For live-imaging of synaptic versus extrasynaptic clustering, synapses were identified by expressing intrabodies against the synaptic marker proteins PSD95 and gephyrin, to simultaneously label excitatory and inhibitory synapses, as we have described recently (Buonarati et al., 2020; Cook et al., 2019); the YFP-tagged CaMKII isoforms were expressed to visualize clustering before and after excitotoxic glutamate insults (100 μM for 5 min). CaMKII clusters were detected extrasynaptically, both basally and after stimulation (Figure 5). For both isoforms, excitotoxic stimulation significantly increased extrasynaptic clustering (Figure 5A,C) and enrichment at excitatory synapses (Figure 5B,C). As previously described for the α isoform (Buonarati et al., 2020), no clustering at inhibitory synapses was observed for CaMKIIβ (Figure S4).

**Figure 5.**
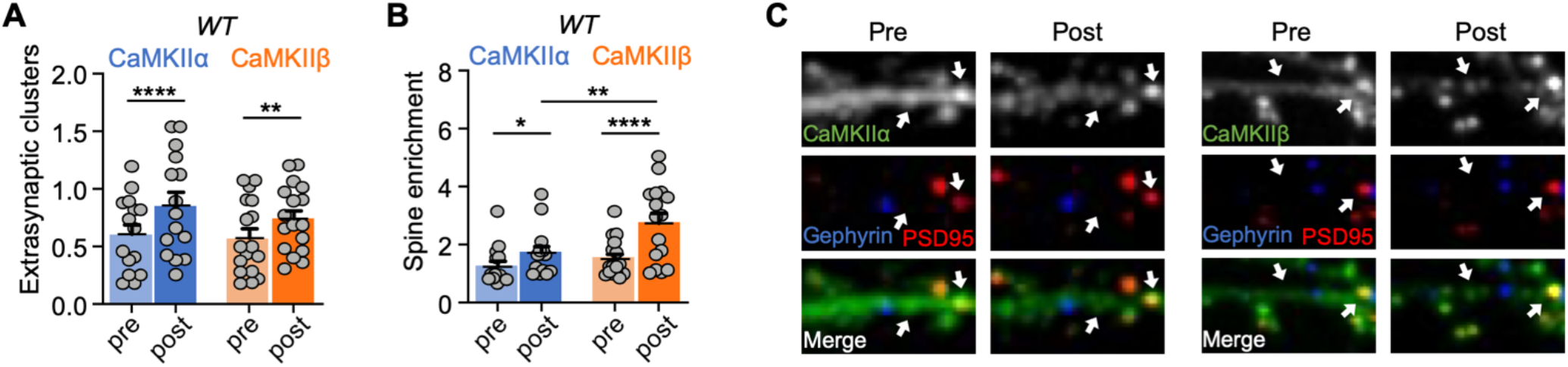
Clustering of CaMKIIα and CaMKIIβ induced by prolonged glutamate in wildtype neurons. Quantifications show mean + SEM. *p<0.05, **p<0.01, ****p<0.0001. (A) Quantification of extra-synaptic clusters induced by excitotoxic glutamate (100 μM glutamate, 5 min) in WT cultured hippocampal neurons (DIV 15-17) (two-way repeated-measures ANOVA, Bonferroni’s test: ****p<0.0001 for CaMKIIα pre vs. post, n=15; **p=0.0049 for CaMKIIβ pre vs. post, n=17). (B) Quantification of excitatory synapse enrichment induced by excitotoxic glutamate (100 μM glutamate, 5 min) in WT cultured hippocampal neurons (DIV 15-17) (two-way repeated-measures ANOVA, Bonferroni’s test: *p=0.0457 for CaMKIIα pre vs. post, n=14; **p<0.0001 for CaMKIIβ pre vs. post, n=16; **p=0.0078 for post CaMKIIα vs. post CaMKIIβ). (C) Representative confocal images show overexpressed CaMKIIα (left) or CaMKIIβ (right), endogenous PSD95 (in red) to mark excitatory synapses, and endogenous gephyrin (in blue) to mark inhibitory synapses.

### Homomeric CaMKIIβ holoenzymes show less propensity for higher order clustering

Somewhat surprisingly, CaMKIIα and β showed the same level of extrasynaptic clustering in hippocampal neurons (Figure 5), even though CaMKIIβ has been described to be incompetent for ischemia-related clustering *in vitro* (Hudmon et al., 2001). Thus, we decided to compare these isoforms also in our *in vitro* clustering assay. As described previously (Vest et al., 2009), ischemic conditions were mimicked by addition of Ca^2+^/CaM and ADP at a low pH of 6.5; then cluster formation was assessed by differential centrifugation. While some amount of both CaMKIIα and β was detected in the 16,000x*g* pellet under basal conditions, this amount dramatically increased under ischemic conditions only for CaMKIIα but not β (Figure 6A,B). By contrast, in 100,000*x g* pellets, both isoforms showed a significant increase in precipitation under ischemic conditions (Fig 6C,D). Nonetheless, the induced precipitation of CaMKIIβ was significantly less than that of CaMKIIα after either centrifugation speed (Figure 6B,D). These results suggest that both isoforms can cluster *in vitro*, but that CaMKIIβ forms less and/or smaller-sized clusters than the a isoform.

**Figure 6.**
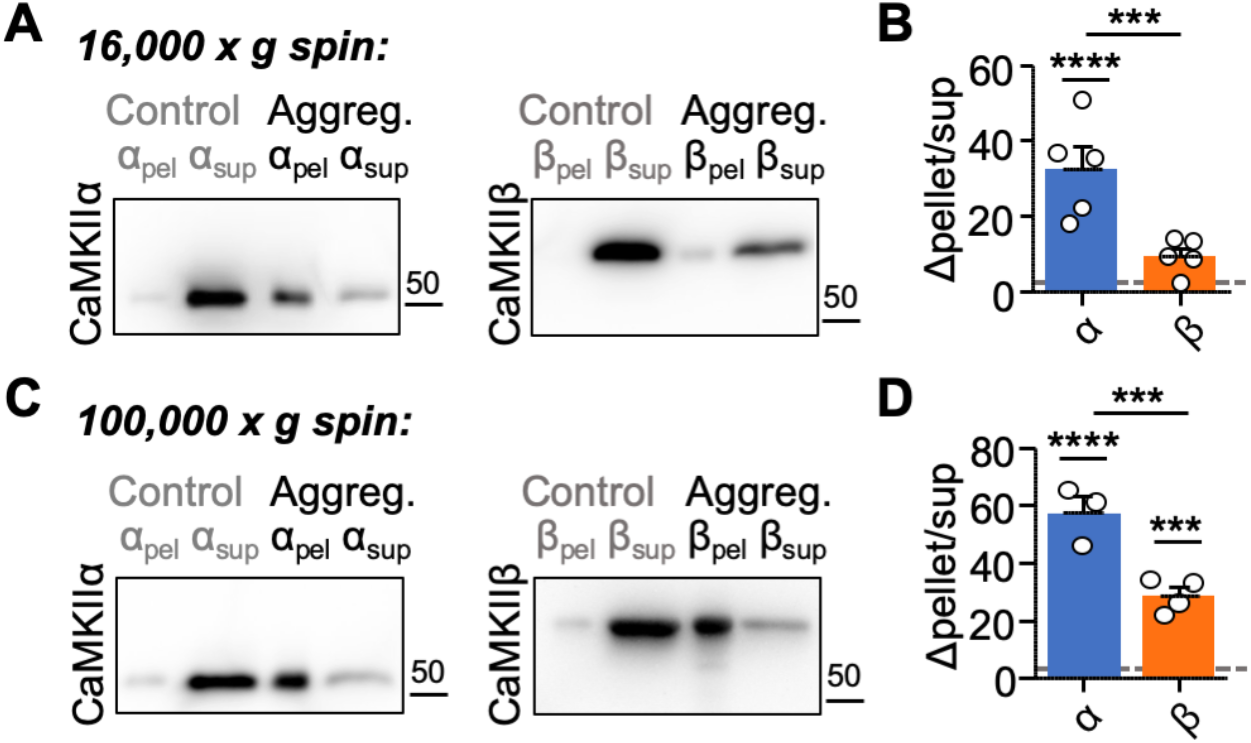
CaMKIIβ self-aggregates less than CaMKIIα *in vitro*. Quantifications show mean ± SEM. ***p<0.001. (A) Representative immunoblots for CaMKIIα and CaMKIIβ. Aggregates were detected in 16,000x*g* pellets following incubation in 2 mM Ca^2+^, 1 μM CaM, and 1 mM ADP at low pH (6.4) for 5 min at room-temperature. Control samples were incubated in 50 mM EGTA at pH 7.4. (B) Quantification of change in pellet enrichment (control samples normalized to 1). Only CaMKIIα showed significant clustering under aggregation conditions, compared to control (two-way ANOVA, Bonferroni’s test: ****p<0.0001 for CaMKIIα). CaMKIIβ shows significantly less self-aggregation compared to CaMKIIα (two-way ANOVA, Bonferroni’s test: ***p=0.0004). (C) Representative immunoblots for CaMKIIα and CaMKIIβ. Aggregates were detected in 100,000x*g* pellets following incubation in 2 mM Ca^2+^, 1 μM CaM, and 1 mM ADP at low pH (6.4) for 5 min at room-temperature. Control samples were incubated in 50 mM EGTA at pH 7.4. (D) Quantification of change in pellet enrichment (control samples normalized to 1). Both CaMKIIα and CaMKIIβ showed significant clustering under aggregation conditions, compared to control (two-way ANOVA, Bonferroni’s test: ****p<0.0001 for CaMKIIα; ***p=0.0004 for CaMKIIβ). Furthermore, CaMKIIβ shows greater self-aggregation in 100,000x*g* pellets compared to 16,000x*g* pellets (two-way ANOVA, Bonferroni’s test: ****p<0.0001). However, CaMKIIβ still shows significantly lower self-aggregation than CaMKIIα even after 100,000x*g* (two-way ANOVA; Bonferroni’s test: ***p=0.0003).

Then why was clustering of CaMKIIα and β indistinguishable in WT neurons? One possibility was that CaMKIIβ might efficiently co-cluster with endogenous CaMKIIα. In order to test this possibility, the clustering experiments were repeated in neurons cultured from CaMKIIα KO mice. In the absence of endogenous CaMKIIα, YFP-CaMKIIβ still clustered both basally and after excitotoxic glutamate insults, but to a significantly lesser extent than YFP-CaMKIIα (Figure 7 and Figure S5). Furthermore, the extrasynaptic CaMKIIβ clustering was significantly lower in the CaMKIIα KO neurons compared to WT neurons (Figure S5H). Together, our *in vitro* experiments with purified protein and our imaging experiments in neurons show that CaMKIIβ can form clusters on its own, but to a significantly lesser extent than the CaMKIIα isoform.

**Figure 7.**
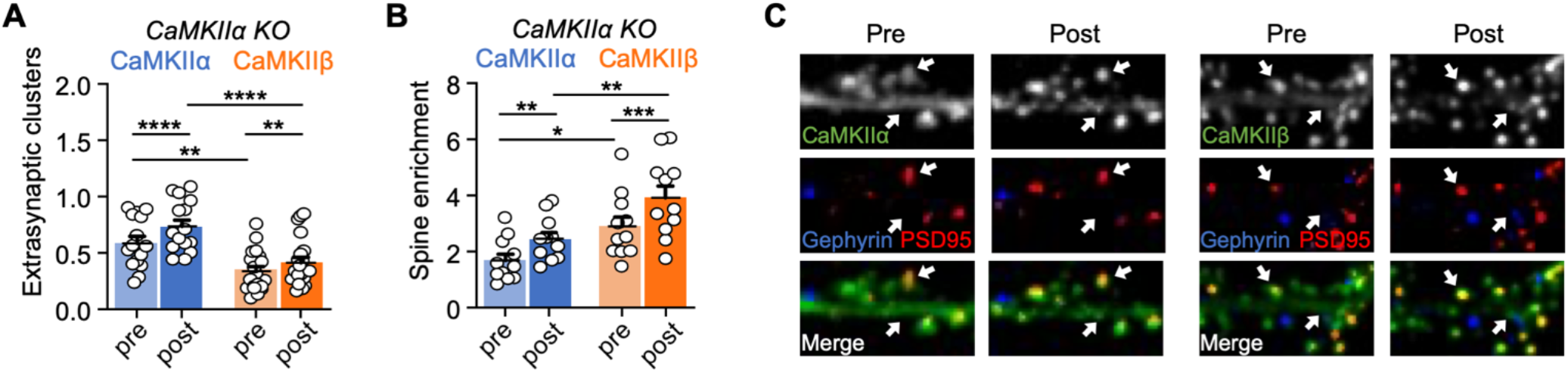
Clustering of CaMKIIα and CaMKIIβ induced by prolonged glutamate in CaMKIIα KO neurons. Error bars indicate SEM in all panels. **p<0.01, ****p<0.0001. (A) Quantification of extra-synaptic clusters induced by excitotoxic glutamate (100 μM glutamate, 5 min) in CaMKIIα KO cultured hippocampal neurons (DIV 15-17) (two-way repeated-measures ANOVA, Bonferroni’s test: ****p<0.0001 for CaMKIIα pre vs. post, n=15; **p=0.0021 for CaMKIIβ pre vs. post, n=20; **p=0.0019 for pre CaMKIIα vs. pre CaMKIIβ; ****p<0.0001 for post CaMKIIα vs. post CaMKIIβ). (B) Quantification of excitatory synapse enrichment induced by excitotoxic glutamate (100 μM glutamate, 5 min) in CaMKIIα KO cultured hippocampal neurons (DIV 15-17) (two-way repeated-measures ANOVA, Bonferroni’s test: **p=0.0025 for CaMKIIα pre vs. post, n=11; ****p<0.0001 for CaMKIIβ pre vs. post, n=11; *p=0.0242 for pre CaMKIIα vs. pre CaMKIIβ **p=0.0051 for post CaMKIIα vs. post CaMKIIβ). (C) Representative confocal images show overexpressed CaMKIIα (left) or CaMKIIβ (right), endogenous PSD95 (in red) to mark excitatory synapses, and endogenous gephyrin (in blue) to mark inhibitory synapses.

## DISCUSSION

Our comparative structure-function analysis of the CaMKIIα and β holoenzymes revealed notable distinctions between these two major neuronal isoforms, including both expected and unexpected structural differences. Perhaps more importantly, it also revealed common structural features of CaMKII that are applicable to the regulation of both isoforms. Specifically, these include a highly dynamic activatable-state conformation, the ability to adopt several oligomeric assemblies (mainly 12-mers, but also 14- and 16-meric holoenzymes) as well as high-order clusters of holoenzymes, and the detection of kinase domain dimer interactions within CaMKIIβ holoenzymes, a mechanism that could mediate the cooperative activation by CaM for all CaMKII isoforms (see Figure 8). As delineated below, these findings provided both answers and new questions.

**Figure 8.**
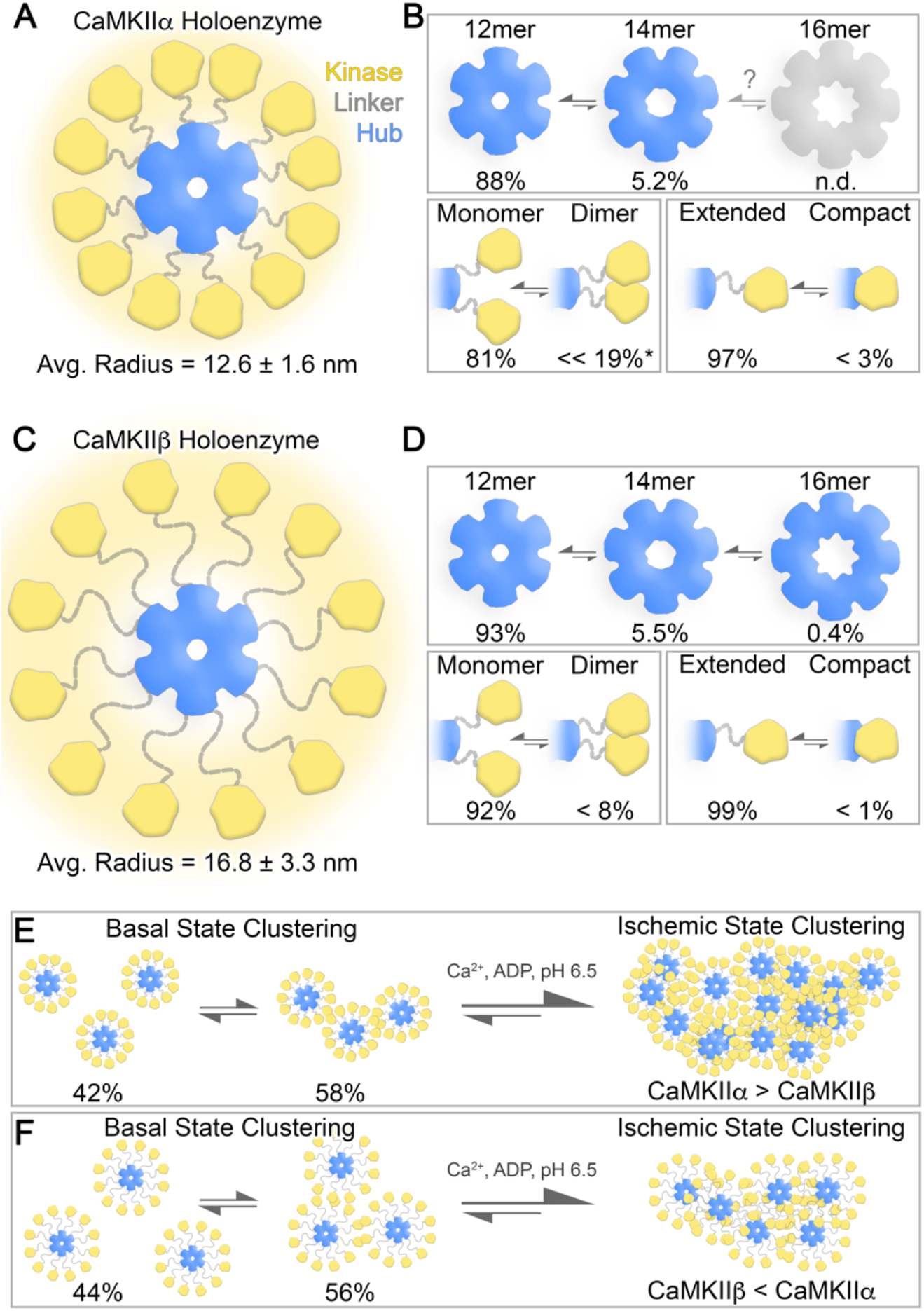
Overview of structural states supported by CaMKII holoenzymes under basal and ischemic conditions. (A) Illustration of the 12-meric CaMKIIα holoenzyme under basal conditions, where a flexible linker region (grey) supports a continuum of extended configurations of kinase domains (yellow) with an average radius of 12.6 ± 1.6 nm (SD), from the center of the hub domain (blue). The halo of yellow density represents the variability in kinase domain positioning observed at the single particle level. (B) *Top*, Oligomeric states of the hub domain resolved by single particle EM, with the 12-mer most populated (88%), followed by the 14-mer (5.2%) and the 16-mer was not detected (n.d.). *Bottom Left*, Kinase domains where predominantly resolved as monomers (81%), with a significant population resolved as putative dimers (<< 19%*). Asterisk indicates that this value is likely over estimated due to artifacts associated with local crowding in the CaMKIIα isoform (see main text). *Bottom Right*, The majority of kinase domains were resolved in an extended state (97%), while a small subset of subunits were localized within contact distance of the hub domain, consistent with a compact state (< 3%). (C) The 12-meric CaMKIIβ holoenzyme, displayed as in panel A, supported a significantly larger extension of kinase domain configurations under basal conditions, as compared to CaMKIIα, with an average radius of 16.8 ± 3.3 nm (SD) facilitated by the longer linker region. (D) *Top*, CaMKIIβ hub domains were resolved predominantly as 12-mers (93%), followed by a 14-mers (5.5%), as well as a novel 16-meric state (0.4%). *Bottom Left*, Kinase domains where also predominantly resolved as monomers (92%), with a significant population resolved as putative dimers (< 8%), (*Bottom Right*) while the compact state is consistent with only < 1% of the population of subunits. (E and F) Illustrate high-order clustering of holoenzymes detected under basal and ischemic conditions for CaMKIIα (E) and CaMKIIβ (F). Under basal conditions, both isoforms form small clusters or pairs of holoenzymes mediated by kinase domain interactions (58% for CaMKIIα and 56% for CaMKIIβ). Under ischemic conditions induced in cells or *in vitro*, there is a shift in the equilibrium toward higher-order clusters, where CaMKIIα clustering was found to be significantly greater than for CaMKIIβ.

### Holoenzyme expansion and autophosphorylation kinetics

The most obvious difference between the CaMKIIα and β holoenzyme was in their radius of expansion, both in their average radius (12.6 vs 16.8 nm) and in their maximal radius (16 vs 23 nm). This difference appears to be also the most predictable one, based on the different lengths of their variable linker regions that tethers the kinase domains to the hub formed by the association domains. Based on the average holoenzyme radii, we calculated a local concentration of kinases domains, within the space occupied by a holoenzyme, to be 2 mM for CaMKIIα and 1 mM for CaMKIIβ. However, there was no apparent difference in the kinetics of the regulatory T286/T287 autophosphorylation that occurs between the subunits of a holoenzyme. While this result seemed counterintuitive at first glance, (i) it is predicted by simple Michaelis-Menten kinetics (Dyla and Kjaergaard, 2020), although (ii) simple Michaelis-Menten kinetics should not necessarily be expected for this autophosphorylation reaction. When T286 is presented as an exogenous substrate on a peptide, its K_m_ is ~10 μM (Coultrap et al., 2010); with this K_m_ and with 2 versus 1 mM substrate concentration, Michaelis-Menten kinetics predicts reaction speeds of 99.50% vs 99.01% of V_max_, *i.e*. a miniscule difference that cannot be resolved in our analyses. However, within the CaMKII holoenzyme, Michaelis-Menten kinetics would break down at least after the first or second autophosphorylation reaction, due to the significant substrate depletion. Additionally, there could have been distinct steric positioning of kinase domains in the CaMKIIα versus β holoenzyme that could either facilitate or reduce the inter-subunit autophosphorylation. Indeed, for the CaMKIIα holoenzymes, there was a discernable favored relative positioning of the kinase domains (Myers et al., 2017), whereas this was not the case for CaMKIIβ. Nonetheless, even for CaMKIIα, the major observation in single particle analysis was the high degree of variability in the kinase domain positioning, consistent with relatively unhindered kinase domain movement and with the similar T286 autophosphorylation kinetics found here for both isoforms.

By contrast, the inhibitory autophosphorylation at T305/306 in CaMKIIα and T306/307 in CaMKIIβ has recently been shown to differ between the isoforms, and this has been attributed to the different linker lengths (Bhattacharyya et al., 2020b). However, whereas T286 autophosphorylation occurs exclusively in *trans* between two subunits of a holoenzyme (Hanson et al., 1994; Rich and Schulman, 1998), the T305/306 autophosphorylation can occur in *cis* within the same subunit, which is then acting both as kinase and as substrate (Colbran, 1993). While such a *cis* phosphorylation reaction should be independent of the concentration of the kinase subunits, it should be more dependent on the steric accessibility of the phosphorylation site and the kinase subunit, as indeed observed (Bhattacharyya et al., 2020b). Similarly, the different linker lengths in the CaMKIIα versus β isoforms may also affect steric accessibility to external substrate proteins, particularly when the holoenzymes are anchored at postsynaptic protein scaffolds.

### Higher order assemblies and subcellular CaMKII targeting

A more surprising difference was the reduced propensity of CaMKIIβ to form higher order clusters under ischemic conditions, both *in vitro* and within neurons. While it has been previously reported that CaMKIIβ lacks excitotoxicity/ischemia-related clustering (Hudmon et al., 2001), the longer linker in CaMKIIβ should instead have been expected to facilitate this clustering: In the a isoform, clustering is thought to be mediated by kinase-domain pairing between holoenzymes (Hudmon et al., 1996; Vest et al., 2009), and a longer linker should facilitate such inter-holoenzyme pairings. The explanation might be that CaMKIIβ can form such pairings, but that the smaller CaMKIIα holoenzymes can pack into larger and/or denser clusters (see Figure 8E,F). Indeed, this notion is supported by the preferential detection of CaMKIIβ *in vitro* clustering by high-versus low-speed centrifugation (a fact that may also explain the previous failure to detect these clusters at all). Furthermore, while extra-synaptic CaMKIIβ clustering was significantly less compared to CaMKIIα, a significant level of clustering was observed also for the β isoform, even in CaMKIIα knockout neurons.

Notably, together with the decrease in extra-synaptic clusters, we observed an increase in synaptic CaMKIIβ clusters. This result appears to be in conflict with the notion that the inter-holoenzyme aggregation mediates clustering not only at extra-synaptic sites but also at synapses (Hudmon et al., 2005), a form of subcellular CaMKII movement that is thought to be important in LTP (Barcomb et al., 2016; Barria and Malinow, 2005; Halt et al., 2012; Incontro et al., 2018; Sanhueza et al., 2011). Indeed, it is well established that CaMKII movement to excitatory synapses requires CaMKII binding to the NMDA-receptor subunit GluN2B (Bayer et al., 2001; Buonarati et al., 2020; Halt et al., 2012). However, this does not rule out the possibility that holoenzyme aggregation contributes to this targeting. Furthermore, the clearly reduced propensity of CaMKIIβ to cluster at extra-synaptic sites does not fully rule out the possibility that its clustering could be enhanced at synaptic sites. For instance, the larger holoenzyme radius and less dense clusters of CaMKIIβ might be favorable within the protein-concentrated environment at postsynaptic densities at excitatory synapses. Nonetheless, our results indicate that the higher-order aggregation of CaMKII holoenzymes plays a more important role in cluster formation at extra-synaptic versus synaptic sites.

The function of extra-synaptic CaMKII clustering is still unclear, but it has been proposed to provide neuro-protection by curbing the over-activation of CaMKII after excitotoxicity/ischemia (Aronowski et al., 1992; Hudmon et al., 1996). This mechanism is clearly insufficient to completely prevent the neuronal cell death after excitotoxic/ischemic insults, but CaMKII inhibition can indeed protect neurons in mouse models of stroke or global cerebral ischemia (Deng et al., 2017; Vest et al., 2010). Further, the CaMKII T286A mutation increases clustering (Hudmon et al., 2005; Vest et al., 2009) and decreases neuronal cell death (Deng et al., 2017). However, this correlation allows only limited conclusions, as the T286A mutation also prevents generation of Ca^2+^-independent autonomous CaMKII activity.

### Multivalent interactions within the holoenzymes and cooperative CaMKII regulation

While kinase domain interactions between holoenzymes are thought to mediate the aggregation of holoenzymes into higher-order assemblies, our results here show the first direct evidence of kinase domain dimer formation also within the holoenzyme. Previous FRET studies have suggested kinase domain dimers are supported by the CaMKII holoenzyme in cells, but direct detection of dimer formation was not resolved in these studies (Nguyen et al., 2015; Nguyen et al., 2012; Thaler et al., 2009). Dimer formation was directly observed in the first crystal structure of a CaMKII kinases domain (specifically for a *C. elegans* CaMKII that was truncated after the regulatory domain; Rosenberg et al., 2005). The low affinity of this interaction (Kd > 100 μM) could be sufficient to support dimer formation based on the 1 – 2 mM concentration of kinase domains within a holoenzyme. Kinase domain dimerization has also been observed biochemically for isolated kinase domains of all four human isoforms, with similarly low-affinity (Kd’s ~200 – 600 μM) (Rellos et al., 2010). However, it is was entirely unclear if such dimer formation would be possible for kinase domains that are tethered to the central hub of association domains within the holoenzyme. The extended state of CaMKIIβ holoenzymes facilitated the ability to directly resolve kinase domain dimers by EM; while this did not allow detailed resolution of the dimerization surface, the defined bi-lobed densities are consistent with the crystallized CaMKII kinase domain dimer structure.

For the smaller CaMKIIα holoenzymes, local crowding of kinase domains limited the ability to clearly resolve kinase domain dimers in CaMKIIα holoenzymes by 2D classification methods, but similar bi-lobed densities could be isolated at the single-particle level (Myers et al., 2017). Thus, similar kinase domain dimers are expected within both CaMKIIα and β holoenzymes. The putative population of kinase domain dimers for CaMKIIα and β were determined to be 19% and 8%, respectively, based on single particle distance measurements obtained from EM micrographs and the average center-to-center distance of ~4.5 nm separating the two subunits in the crystal structure of the kinase domain dimer (see Fig. 3E). Notably, the distribution of CaMKIIβ kinase separation distances are weighted more heavily toward shorter separation distances, and using a slightly larger threshold of 5.0 nm for kinase domain dimerization (the difference of approximately one pixel in the micrographs) would indicate a population of ~16% dimers for the β holoenzymes (and up to ~30% for α holoenzymes). Additional limitations of this indirect approach should also be noted. For example, the higher value for the CaMKIIα holoenzymes is likely an overestimate. This is because the shorter linker in CaMKIIα holoenzymes is expected to result in a higher chance of randomly positioning two kinase domains in close spatial proximity when deposited onto the EM grids (asterisk in Figure 8D). Thus, the dimer population in these two isoforms may be more similar than our reported values indicate, with a likely range of ~8 – 20% kinase domain dimers for both isoforms.

If CaMKIIα and β holoenzymes contain similar fractions of dimerized kinase domains, both isoforms should show a similar level of cooperative activation by CaM, as was indeed observed here. The dimers are proposed to generate cooperativity, as they are formed via interactions of the CaM-binding regulatory domains (Rosenberg et al., 2005). Thus, when CaM binds to one subunit, it would disrupt the dimer and thereby also facilitate binding to the other subunit. While this kinase domain dimerization predicts the observed cooperativity, it also raises some questions. How could a relatively minor fraction of dimers (*e.g*., ~8 – 20%) cause cooperativity with a relatively large Hill coefficient of 1.6?

Our results also suggest the possibility that kinase domain dimers between holoenzymes might contribute to the cooperative activation. As discussed above, these kinase domain interactions are thought to mediate the formation of large holoenzyme aggregates under ischemic conditions. Under basal conditions, only a small fraction of CaMKII precipitated *in vitro* even during high-speed centrifugation. However, some clusters formed in neurons also basally, and small basal assemblies of only a few holoenzymes (which were indeed observed on the EM grid) would be expected to remain soluble even in high-speed centrifugation. Thus, a functional relevance of such small higher order holoenzyme assemblies should not be ruled out, including for regulation of kinase activity. However, it is not clear if the inter-holoenzyme kinase domain interactions can occur by similar mechanism as the kinase domain dimers within holoenzymes; in fact, the low affinity of the dimers that form within holoenzymes makes it unlikely that the same interaction would occur between holoenzymes.

An additional, or alternative, proposed mechanism for the cooperativity lies in a compact conformation in which a kinase domain folds back onto its own association domain (Chao et al., 2011). A recent elegant cryo-EM study indeed directly demonstrated the existence of this conformation (Sloutsky et al., 2020), whereas our previous and current studies could only set an upper limit to its prevalence (Myers et al., 2017). However, with less than 3% of subunits in the compact state, this maximal prevalence is extremely low and consistent to all of the studies (Myers et al., 2017; Sloutsky et al., 2020), which makes it an even less likely candidate mechanism for the observed cooperativity than the kinase domain dimers. Moreover, while the compact conformation should decrease CaM affinity, the structural basis for how it would induce cooperativity is unclear. Such a mechanism would require that the compact conformation of one subunit would affect the positioning of its neighboring subunits. Indeed, kinase domain positioning (as determined by the length of the variable linker region) affects CaM binding: CaMKIIβ is more readily activated by lower concentrations of CaM than the CaMKIIα isoform that has a shorter linker region (Brocke et al., 1999; De Koninck and Schulman, 1998) as also observed in this study. Additionally, a similar relationship is seen also between CaMKIIβ and its shorter linker splice variant CaMKIIβe’ (Bayer et al., 2002; Sloutsky et al., 2020), *i.e*. between two CaMKIIβ variants that share the exact same CaM binding region. However, if any, a compact conformation would appear more likely to promote a more extended conformation of a neighbor, which would already basally facilitate the activation of this neighbor by CaM and would thus be the opposite effect as would be required for cooperativity.

Then, a more likely “neighbor effect” may be that kinase domain dimerization (or disruption of the dimer) affects CaM binding not only to the dimer itself, but also to the neighboring subunits. In this way, one single kinase domain dimer could potentiate CaM binding to half of the subunits within a 12-meric CaMKII holoenzyme, *i.e*. the dimer pair itself plus its four neighbors. A similar model was proposed previously, but assumed that the neighbors would also be dimers (Chao et al., 2011); however, with the observed lower occurrence of dimers, the model would have to be modified to include effects also on non-dimerized neighbors. Indeed, such a model is consistent with the emerging view of CaMKII holoenzyme structure, one that is not constrained by a single defined state, but rather a highly dynamic conformational ensemble characterized by multiple transient low-affinity interactions (see Figure 8).

An intriguing comparison can be made to other systems hallmarked by multivalent low-affinity interactions organized by intrinsically disordered protein domains that are capable of forming biomolecular condensates (Brangwynne et al., 2009). We suggest that the unique structural and biophysical properties of the CaMKII holoenzyme structure (*e.g*., high local concentration of multivalent binding modes) may facilitate the formation of a molecular-scale condensate, at least from a conceptual point of view. Condensates (*i.e*., liquid-liquid phase separation or LLPS) have emerged as a novel regulatory mechanism at synapses (Chen et al., 2020) and the CaMKII interaction with GluN2B has recently been shown to support condensate formation (Hosokawa et al., 2020). What is intriguing about this analogy of individual CaMKII holoenzymes to condensates is that the dissolution of biomolecular condensates is highly cooperative (Banani et al., 2017; Li et al., 2012). Such a model could explain how activation of CaMKII holoenzymes can achieve a higher degree of cooperativity than would be expected based on the extent of kinase domain dimers. Notably, some studies have described Hill coefficients for CaMKII activation by CaM that are even higher than the Hill coefficient of ~1.6 reported here (Chao et al., 2011; Rosenberg et al., 2005). In a model analogous to molecular condensates, each kinase domain exists in an equilibrium between rapidly exchanging interactions involving multiple neighboring kinase or hub domains, a notion supported by the flexibility of kinase domain positioning with the holoenzymes. The activation of one kinase domain would then disrupt the interaction with multiple neighboring kinase domains, leading to the cooperative collapse or dissolution of the basal-state. While there is currently no direct evidence that a kinase domain dimer (or kinase-hub complex) can affect the positioning of neighboring kinase domains within the holoenzyme, such a proposition appears to be at least more plausible than what can be explained by any single-defined state of CaMKII.

### Beyond the 12-mer: Outlook for future studies

The oligomeric state of CaMKII holoenzyme is clearly important for facilitating its physiological roles Ca^2+^-frequency detection and in regulating LTP and LDP (Coultrap et al., 2014; De Koninck and Schulman, 1998; Giese et al., 1998; Hanson et al., 1994). The predominant state of both CaMKIIα and β holoenzymes is the 12-meric assembly (see Figure 2 and Myers et al., 2017). However, both holoenzymes can also support 14-meric assemblies, and here we show that CaMKIIβ can even support a 16-meric assembly. To our knowledge, this is the first time such a high oligomeric state of the CaMKII hub domain has been detected in metazoans. Interestingly, bacteria and algae species contain orphan proteins with sequence and structural homology to CaMKII hub domains, that adopt 16-to 20-mers (McSpadden et al., 2019). Crystallographic analysis of the hub-like assembly from *Chlamydomonas reinhardtii* revealed an 18-meric structure with striking similarity to CaMKII hub assemblies, but with increased hydrogen-bonding at the lateral subunit interface. Remarkably, when these hydrogen-bonding residues were incorporated into the CaMKIIα hub domain it assembled as 14- and 16-mers (McSpadden et al., 2019). It is currently unclear if wildtype CaMKIIα holoenzymes support a 16-meric assembly, but if it can it is likely a very minor population as we have found that this state only represents ~0.4% of the population in CaMKIIβ holoenzymes under basal-conditions, and both isoforms lack the hydrogen-bonding potential identified in the algae hub-like assembly.

These observations raise the important question as to what is the functional significance of higher-order oligomeric states of the CaMKII holoenzyme? It has recently been shown that under activating conditions, CaMKII subunits (dimeric pairs) are capable of undergoing exchange between other activated or non-activated holoenzymes (Bhattacharyya et al., 2016; Stratton et al., 2014). While subunit exchange was minimal under basal conditions, it is possible that the small population of 14-mer/16-mers represented high-energy intermediate states involved in the subunit exchange mechanism. Activation of the CaMKII holoenzyme is thought to destabilize the hub complex, through interactions with the regulatory domain (Bhattacharyya et al., 2016; Karandur et al., 2020; Stratton et al., 2014). Thus, the 14-mer and 16-mer states could also provide a storage mechanism for pools of potentiated subunits to be released under activating conditions. Intriguingly, this subunit-exchange mechanism has been proposed to enable the propagation of CaMKII activation and other neuronal plasticity mechanisms that could play important roles in learning and memory (Bayer and Schulman, 2019; Bhattacharyya et al., 2020a). Future studies will be needed to further test these hypotheses, and to shed light on the mechanistic basis for how such regulatory functions are achieved.

## Materials and methods

### Key resources table

| Reagent type (species) or resource | Designation | Source or reference | Identifiers | Additional information |
| --- | --- | --- | --- | --- |
| Strain (*Mus musculus*), C57BL/6J | Wild type | Charles River Laboratory |  | Primary hippocampal cultures |
| Strain (*Mus musculus*), C57BL/6J | CaMKIIα KO | (Coultrap et al., 2014) | | Primary hippocampal cultures |
| Strain (*E. coli*) | One Shot BL21 star bacteria | Invitrogen | C601003 | Subcloning |
| Recombinant DNA reagent | PSD-95-FingR-GFP | (Gross et al., 2013) | Addgene_46295 |  |
| Recombinant DNA reagent | Gephyrin-FingR-GFP | (Gross et al., 2013) | Addgene_46296 |  |
| Recombinant DNA reagent | YFP-CaMKIIα | (Bayer et al., 2006; Shen and Meyer, 1999) | | |
| Recombinant DNA reagent | YFP-CaMKIIβ | Made in house | | |
| Antibody | anti-CaMKII pan | Genetex | GTX127939 | 1:1000 |
| Antibody | anti-CaMKIIα | Made in house | CBα2 | 1:1000 |
| Antibody | anti-CaMKIIβ | Made in house | CBβ1 | 1:1000 |
| Antibody | anti-pT286 | Phospho-Solutions | p1005-286; RRID: AB_2492051 | 1:2000 |
| Antibody | anti-Rabbit | GE Healthcare | NA934V; RRID: AB_772206 | 1:10,000 |
| Antibody | anti-Mouse | GE Healthcare | NA931V; RRID: AB_772210 | 1:10,000 |
| Chemical compound, drug | Glutamate | Sigma | 6106-04-3 | Excitotoxic stimuli in neurons |
| Chemical compound, drug | Papain | Worthington | LS 03126 | Dissociating neurons |
| Chemical compound, drug | Lipofectamine 2000 | Invitrogen | 11668027 | Coating glass coverslips |
| Chemical compound, drug | B-27 supplement | GIBCO | 17504044 | Neuronal media supplement |
| Chemical compound, drug | cOmplete protease inhibitor cocktail | Roche | 1187380001 | Protease inhibitor |
| Chemical compound, drug | Microcystin-LR | Calbiochem | 475815 | Phosphatase inhibitor |
| Chemical compound, drug | Uranyl formate | SPI-Chem | 16984-59-1 | Electron microscopy |
| Chemical compound, drug | Sodium hydroxide | Fisher Scientific | 1310-73-2,497-19-8 | Electron microscopy |
| Other | 400 mesh, 3.0 mm O.D. copper grid | PELCO | 1GC400 | Electron microscopy |
| Peptide, recombinant protein | Calmodulin | Made in house | CaM | Activation of CaMKII |
| Peptide, recombinant protein | Ca<sup>2+</sup>/CaM-dependent kinase II $\alpha$ | Made in house | CaMKIIα | Purified kinase |
| Peptide, recombinant protein | Ca<sup>2+</sup>/CaM-dependent kinase II $\beta$ | Made in house | CaMKIIβ | Purified kinase |
| Critical commercial assay | Pierce BCA protein assay | Thermo Fisher | 23225 | Measuring protein |
| Critical commercial assay | SuperSignal West Femto | Thermo Fisher | 34095 | Developing immunoblots |
| Software, algorithm | Slidebook 6.0 | Intelligent Imaging Innovations (3i) | RRID:SCR_014300 |  |
| Software, algorithm | Prism 7.0 | Graphpad | RRID:SCR_002798 |  |
| Software, algorithm | ImageJ | NIH | RRID:SCR_003070 |  |
| Software, algorithm | EMAN2.1 | NIH | RRID:SCR_018867 |  |
| Software, algorithm | RELION3.0b | MRC (United Kingdom) | RRID:SCR_016274 |  |
| Software, algorithm | UCSF Chimera | NIH | RRID:SCR_004097 |  |
| Software, algorithm | Python 3 | Python Software Foundation | RRID:SCR_008394 |  |
| Software, algorithm | Microsoft Excel | Microsoft | RRID:SCR_016137 |  |

### Animals

All animal procedures were approved by the Institutional Animal Care and Use Committee (IACUC) of the University of Colorado Anschutz Medical Campus and were carried out in accordance with NIH best practices for animal use. All animals were housed in ventilated cages on a 12 h light/ 12 h dark cycle and were provided ad libitum access to food and water. Mixed sex pups from homozygous mice (P1-2; on a C57BL/6 background) were used to prepare dissociated hippocampal cultures for live imaging. The CaMKII KO mice were described previously (Coultrap et al., 2014).

### DNA constructs

CaMKIIα and β were subcloned into a YFP expression vector (pcDNA3 backbone; Addgene #13033). Expression vectors for the GFP-labeled FingR intrabodies targeting PSD-95 and gephyrin were kindly provided by Dr. Donald Arnold (University of Southern California, Los Angeles, CA, USA) as previously characterized (Gross et al., 2013; Mora et al., 2013). The fluorophore label was exchanged using Gibson Assembly to contain the following tags in place of GFP: PSD-95-FingR-mTurquois and gephyrin-FingR-mCherry.

### CaMKII and CaM purification

For biochemistry and electron microscopy, homomeric rodent CaMKIIα and β was purified after expression in S*f*9 cells, and CaM was purified after expression in bacteria as previously described (Coultrap and Bayer, 2012a; Singla et al., 2001).

### Electron microscopy

Full length CaMKIIβ holoenzymes purified from eukaryotic S*f*9 cell expression were prepared for negative stain EM by diluting a freshly thawed aliquot of protein (1:40 vol vol^-1^) in EM buffer containing 50 mM HEPES (pH 7.4), 120 mM KCl and 0.5 mM EGTA. A 3 μL drop of the diluted specimen (~35 μM) was applied to a glow-discharged continuous carbon coated EM grid (Ted Pella). Excess protein was removed by blotting with filter paper and washing twice with EM buffer. The specimen was then stained with freshly prepared (0.75% wt vol^-1^) uranyl formate (SPI-Chem), blotted and dried with laminar air flow.

Negatively stained specimens were visualized on a 120kV TEM (Tecnai iCorr, FEI) and digital micrographs were manually collected on a 2K x 2K CCD camera (Eagle, FEI) at a nominal magnification of 49,000 x at the specimen level. Micrographs were collected with a calibrated pixel size of 4.37 Å and defocus of 1.5 – 2.5 μM. A total of 907 micrographs were collected and screened for astigmatism and drift based on Thon rings in the power spectra after determination of contrast transfer function (CTF) parameters in EMAN2.1 (Tang et al., 2007). 17,347 single particles images were manually picked in EMAN 2.1 and extracted with a box size of 144 x 144 pixels. Boxed images used for single particle analysis were of isolated holoenzymes that were not associated with neighboring holoenzymes (such as the particles examined for clustering). Reference-free 2D classification and variance analysis was conducted in RELION3.0b (Zivanov et al., 2018) on CTF-corrected (phase-flipped), using various masking strategies: holoenzyme = 550 Å mask, hub domain = 150 Å mask, and kinase domains = inner mask 135 Å, without applied symmetry. For comparative analysis, the previously acquired negative stain EM dataset of 10,902 untilted CaMKIIα single-particle images (Myers et al., 2017) was reprocessed in EMAN 2.1 and RELION3.0b, as described above using a box size of 128 x 128 pixels.

### Single particle measurements and statistical analysis

Statistical analyses of individual holoenzyme particle dimensions were obtained from particle lengths using the measurement tool in EMAN2.1, as previously described (Myers et al., 2017). A radius of extension for individual kinase subunits (*n* = 926 measurements obtained from 95 holoenzymes) was determined by measuring the distance from the center of the pore in the hub domain complex to the center of each peripheral density corresponding to the kinase domains. For each of these measurements, a distance of 22.5 Å was appended (corresponding to the average radius of the kinase domain) to yield a value that represents the full extension of the kinase domain. Inter-molecular kinase separation was determined by measuring from the center of one peripheral kinase density to the center of the nearest clockwise neighboring kinase density. For the linker extension analysis, the hub average radius (55 Å) and the kinase average radius (22.5 Å) was subtracted from the raw hub to kinase radius measurements, as a representative distance of linker extension and comparison to a 2D random walk model 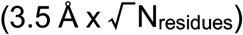 (Flory, 1975). The local concentration of kinase domains in the context of a single dodecameric holoenzyme was determined by assuming a spherical volume with the radius corresponding to the average radius extension for each isoform ±2 x standard deviation appended. To analyze intermolecular clustering of holoenzymes, a random subset of raw micrographs was visually inspected for CaMKIIα (1207 particles) and CaMKIIβ (1255 particles) by counting the total number of holoenzymes and the number of non-clustering holoenzymes (designated as being separated by at least ~1.5 times the particle diameter from a nearest neighbor). For statistical comparisons, an F-test was performed to determine the difference in sample variances, followed by a T-test (two sample assuming unequal variances). All statistical analysis and graphical interpretations were done using libraries in python 3.

### Pseudo-atomic modeling of the CaMKII 16-meric hub assembly and back-projection analysis

A pseudo-atomic model of the hexadecameric (16-mer) CaMKII hub assembly was built manually, using the atomic coordinates corresponding to a vertical hub dimer extracted from the previously published dodecameric assembly (PDB 5IG3) (Bhattacharyya et al., 2016), with an applied C8 symmetry (overall D8 symmetry) that was adjusted to approximately fit within the experimental 2D class average density following C8 symmetry averaging (outer diameter of ~13 nm) and to obtain minimal steric overlap between neighboring hub domains. For comparing with 2D class averages, hub models where filtered to 30 Å and 2D back projections were generated in RELION (Scheres, 2012).

### Hippocampal cultured neurons

To prepare primary hippocampal neurons from WT or mutant mice, hippocampi were dissected from mixed sex mouse pups (P1-2), dissociated in papain for 30 min, and plated at 200-300,000 cells/mL for imaging. At DIV 14-15, neurons were transfected with 1 μg total cDNA per well using Lipofectamine 2000 (Invitrogen). At DIV 16-17, neurons were treated and imaged.

### Live imaging of hippocampal cultured neurons

All images were acquired using an Axio Observer microscope (Carl Zeiss) fitted with a 63x Plan-Apo/1.4 numerical aperture objective, using 445, 515, 561, and 647 nm laser excitation and a CSU-XI spinning disk confocal scan head (Yokogawa) coupled to an Evolve 512 EM-CCD camera (Photometrics). During image acquisition, neurons were maintained at 34°C in 10 mM HEPES-buffered neuronal media. After baseline imaging (‘pre’), cells were treated with 100 μM glutamate and imaged 5 min later (‘post’). Tertiary dendrites from pyramidal spiny neurons were selected from maximum intensity projections of confocal Z stacks. Slidebook 6.0 software (Intelligent Imaging Innovations [3i]) was used to analyze CaMKII-YFP at excitatory (PSD-95) and inhibitory (gephyrin) synapses. Specifically, the mean YFP intensity at PSD-95 or gephyrin threshold masks on a given dendrite was divided by the mean YFP intensity of a line drawn in the adjacent dendritic shaft. ImageJ (National Institute of Health) was used to analyze CaMKII at extra-synaptic sites. Specifically, the thresholded mask for PSD-95 puncta was subtracted from the CaMKII channel and the remaining CaMKII clusters (0.1 μm<cluster<1.0 μm) were quantified as number per 10 μm dendrite.

### CaMKII *in vitro* reactions

CaMKII self-associations assays were performed similar to previous work (Vest et al., 2009). Purified CaMKII (0.5 μM) was pre-cleared by ultracentrifugation (100,000 x g) at 4 °C for 45 min, then combined with 25 μM PIPES pH 6.4, 20 mM KCl, 10 mM MgCl_2_, 0.1 mg/mL bovine serum albumin, 0.1% Tween 20, 0.5 mM dithiothreitol, 2 mM CaCl_2_, 1 μM CaM, and 1 mM ADP. Control samples were instead combined with 25 μM PIPES pH 7.4, 20 mM KCl, 10 mM MgCl_2_, 0.1 mg/mL bovine serum albumin, 0.1% Tween 20, 0.5 mM dithiothreitol, and 50 mM EGTA. The mixtures were prepared on ice and then incubated for 5 min at room temperature prior to centrifugation (16,000 x g) or ultracentrifugation (100,000 x g) at 4 °C for 30 min. CaMKII in the supernatant and pellet was detected via immunoblot.

For autophosphorylation assays, purified CaMKII (0.1 uM) was pre-cleared by ultracentrifugation (100,000 x g) at 4 °C for 45 min, then combined with 25 μM PIPES pH 7.1, 10 mM MgCl_2_, 0.1 mg/mL bovine serum albumin, 4 mM CaCl_2_, 3 μM CaM, and 1 mM ATP. The mixtures were prepared on ice and then incubated at 30 °C for 0 sec, 30 sec, 180 sec, or 15 min. Autophosphorylation at pT286-CaMKIIα and pT287-CaMKIIβ was detected via immunoblot.

For kinase activity assays, purified CaMKII (2.5 nM) was combined with 50 mM PIPES pH 7.1, 10 mM MgCl_2_, 0.1 mg/mL bovine serum albumin, 1 mM CaCl_2_, 100 μM [γ-^32^P] ATP, 75 μM syntide-2 subtrate peptide, and 0.6 nM to 6 μM CaM. The mixtures were prepared on ice and then pre-incubated at 30 °C for 5 min. CaMKII was added and the reaction was allowed to progress for 1 min at 30 °C. To stop the reaction, 40 μL of the 50 μL reaction mixture was spotted onto P81 cation exchange chromatography paper (Whatman) squares. After extensive washes with water, phosphorylation of the substrate peptide bound to the P81 paper was measured by liquid scintillation counting. CaMKII activity (V_m_; reactions/kinase/min) was quantified as a fraction of maximal activity (V_max_) for each experiment. Data were fitted using a non-linear regression with variable slope.

### SDS-PAGE and immunoblot

Protein content was determined using the Pierce BCA protein assay (Thermo Fisher). 4-10 μg of total protein in Laemmli sample buffer was resolved by SDS-PAGE on 9% polyacrylamide gels and transferred to polyvinylidene fluoride (PVDF) membrane at 24 V for 1-2 h at 4°C in transfer buffer containing: 12% MeOH, 25 mM Tris, and 192 mM glycine. All membranes were blocked in 5% nonfat dried milk in TBS-T (20 mM Tris pH 7.4, 150 mM NaCl, 0.1% Tween 20) before primary antibody incubation for 2 h at room temperature or overnight at 4°C. Antibodies and dilutions were as follows: rabbit anti-CaMKII (Genetex; 1:1000), mouse anti-CaMKIIα (in house CBα2; 1:1000), mouse anti-CaMKIIβ (in house CBβ1; 1:1000), and anti-pT286 (Phospho-Solutions; 1:2000). Blots were then washed in TBS-T, incubated in horseradish peroxidase-labeled goat anti-rabbit or goat anti-mouse antibodies (GE Healthcare; 1:10,000) for 1 h at room temperature, and washed again in TBS-T. Immunoreactive signal was visualized by chemiluminescence (Super Signal West Femto, Thermo Fisher) using the Chemi-Imager 4400 system (Alpha-Innotech). Densitometry analysis was performed using ImageJ (National Institute of Health), with all samples normalized to control conditions on the same gel.

### Quantification and statistical analysis

Structural measurement data are shown as mean ± SEM, with standard deviation or 95% confidence interval where indicated using Microsoft Excel or SciPy. Comparisons between isoforms for kinase radius and separation measurements were done with a f-test to determine sample variance differences, followed by a t-test (two sampled assuming unequal variance). Functional data are shown as mean ± SEM and analyzed using Prism (GraphPad) software. Comparisons between pre- and post-glutamate images in neurons were analyzed using repeated measures two-way ANOVA with Bonferroni’s post-hoc test. Self-association assays with purified CaMKII were analyzed using two-way ANOVA with Bonferroni’s post-hoc test. Kinase activity assays to asses CaM dose/response were analyzed using extra-sum-of-squares F-test. Comparisons in WT neurons at inhibitory synapses were analyzed using paired, two-tailed Student’s t-test. Statistical significance and sample size (n) are indicated in the figure legends.

## ACKNOWLEDGEMENTS

We thank Ms. Janna Mize-Berge for help with mouse colony maintenance as well as Mr. Jonathan Flores and Dr. Janette Myers for help with EM grid preparation and the OHSU Multiscale Microscopy Core for instrumentation access and training. The research was funded by National Institutes of Health grants F32AG066536 (to O.R.B.), P30NS048154 (UCD neuroscience center grant), R01NS081248 and R01NS110383 (to K.U.B.), and R35GM124779 and R01EY030987 (to S.L.R.).

## AUTHOR CONTRIBUTIONS

O.R.B., A.P.M., S.J.C., and S.L.R. performed experiments; K.U.B. and S.L.R. conceived this study, with contribution from all authors; K.U.B. and S.L.R. wrote the first draft and all authors contributed to the final manuscript.

## SUPPLEMENTAL FIGURES AND LEGENDS

**Figure S1.**
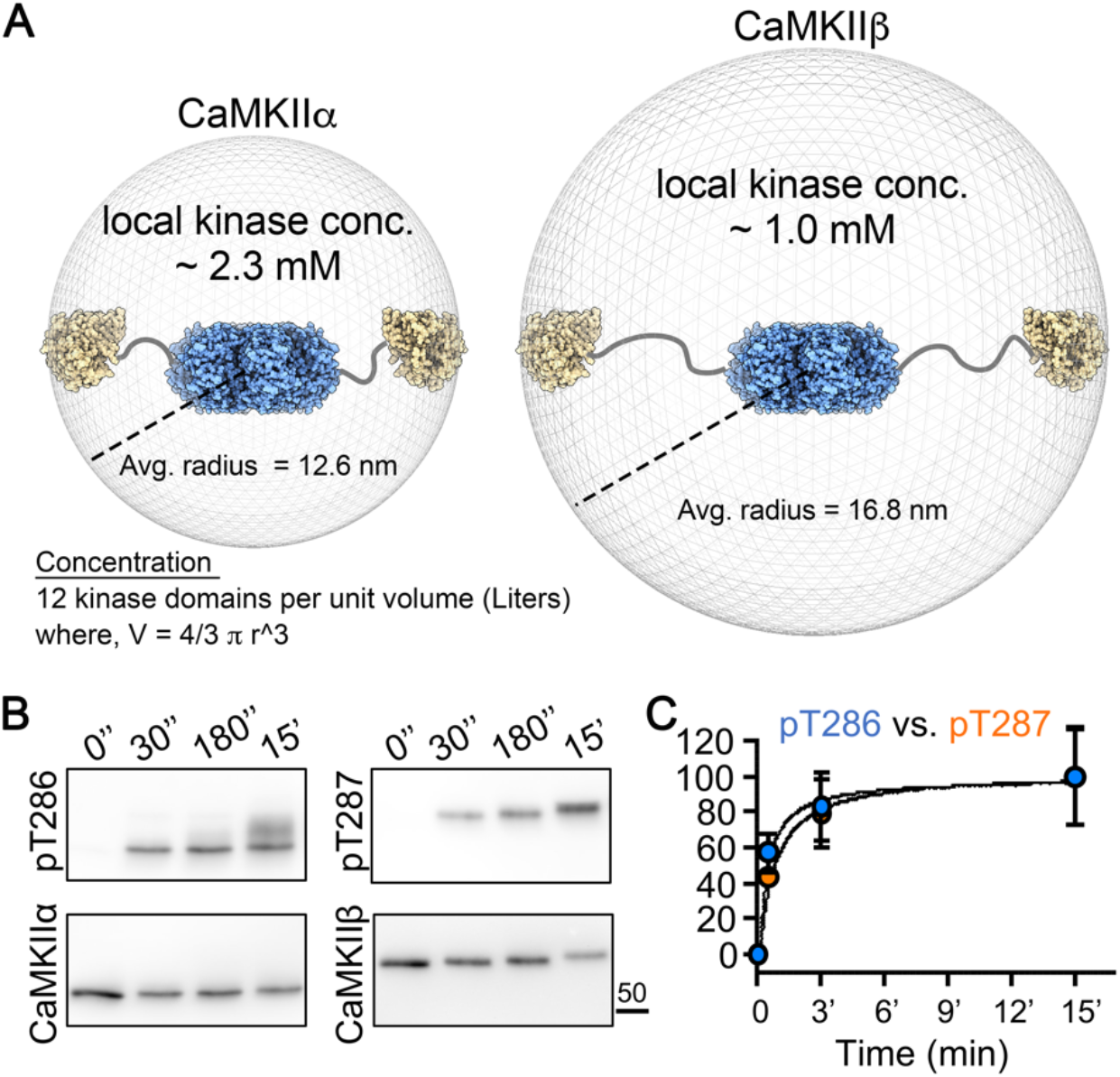
Effect of local kinase domain concentration on pT286-CaMKIIα and pT287-CaMKIIβ in vitro kinetics. Quantifications show mean ± SEM. (A) Estimated local kinase concentration in CaMKIIα (*left*) and CaMKIIβ (*right*) holoenzymes. Calculations assume individual kinase domains (12 per holoenzyme) may occupy a spherical volume that is approximated by the average radius of the holoenzyme complex. (B) Representative immunoblots showing autophosphorylated T286-CaMKIIα (*left*) and T287-CaMKIIβ (*right*) after 0 sec, 30 sec, 180 sec, and 15 min at 30° C. Reactions were performed with 100 nM purified kinase in buffered solution containing 2 mM Ca^2+^, 3 μM CaM, 10 mM Mg^2+^ and 1 mM ATP. (C) Quantified time-course of normalized pT286-CaMKIIα versus pT287-CaMKIIβ (n=4). Solid lines through the data points represent nonlinear best-fits.

**Figure S2.**
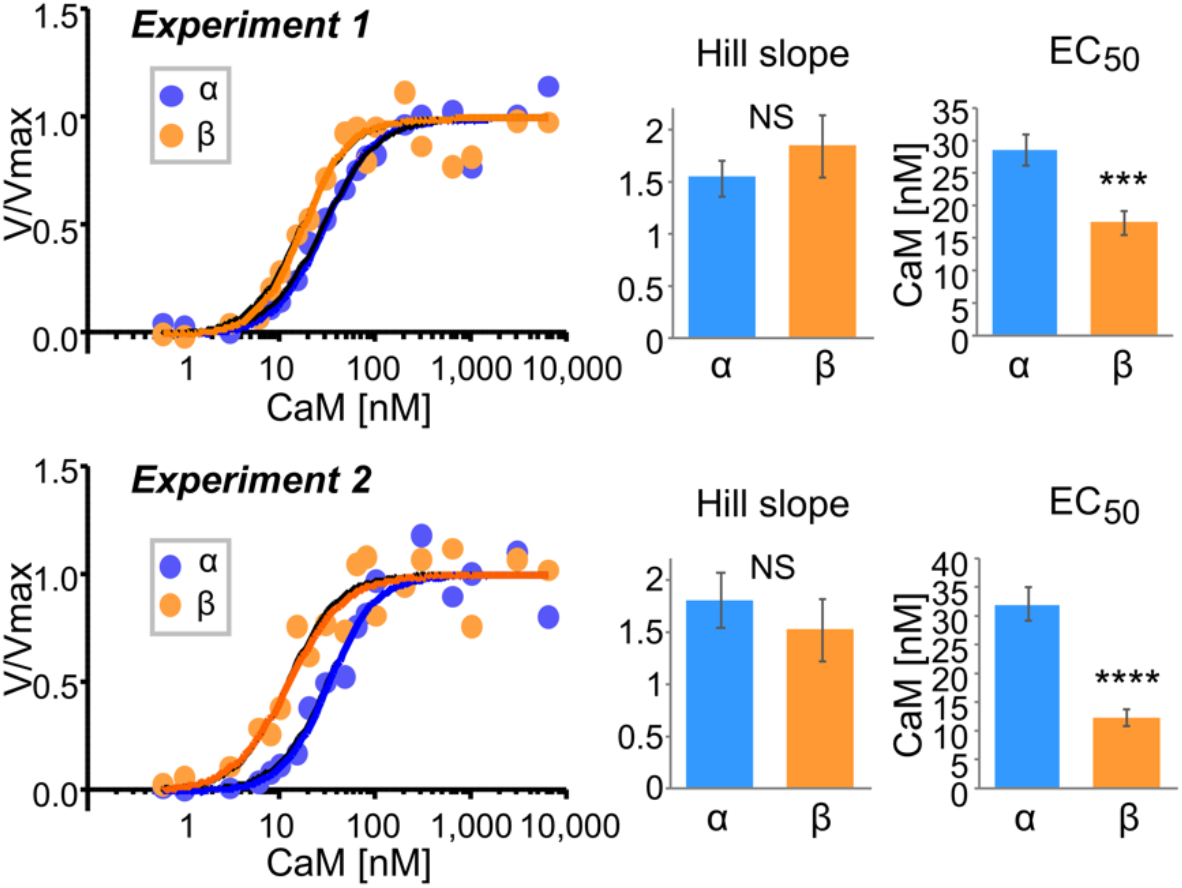
CaMKIIα and CaMKIIβ activation by CaM: individual experiments. Quantifications show mean ± SEM. ***p<0.001, ****p<0.0001. *In vitro* CaMKII activity in response to varying Ca^2+^/CaM (0.6 nM to 6 μM CaM). The two curve fits shown are based on data from independent reactions: experiment 1 (*top*) and experiment 2 (*bottom*). Both experiments demonstrated a Hill slope >1 for CaMKII, with no differences detected between α and β isoforms (extra-sum-of-squares F-test, p=0.3619 for experiment 1 and p=0.4921 for experiment 2). However, both experiments showed a reduced EC_50_ for CaMKIIβ compared to α (extra-sum-of-squares F-test, ***p=0.006 for experiment 1 and ****p<0.0001 for experiment 2).

**Figure S3.**
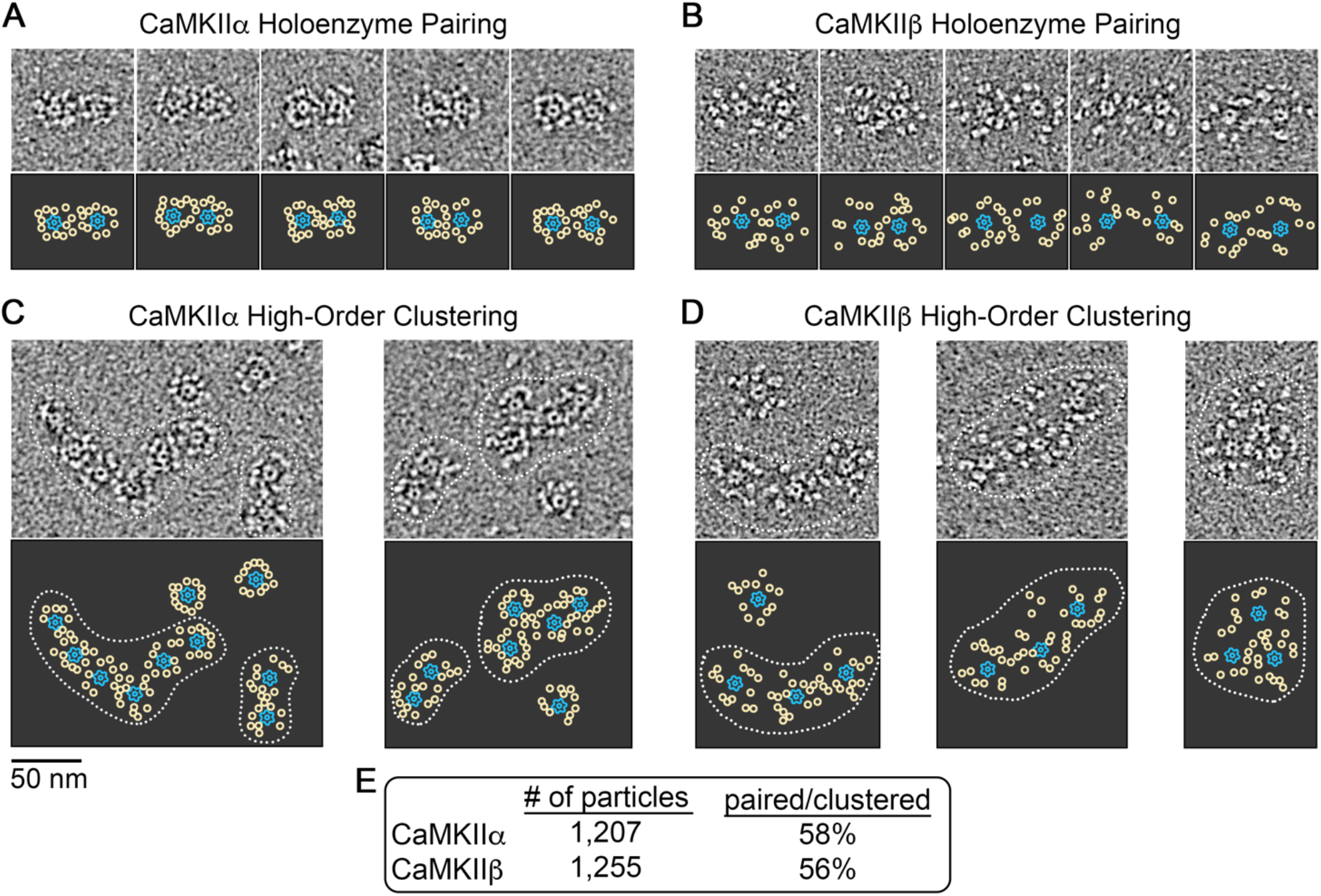
EM analysis of CaMKII holoenzyme clustering under basal conditions. (A and B) *Top*, representative EM images showing apparent holoenzyme paring for CaMKIIα and CaMKIIβ, respectively. *Bottom*, displays annotated representation of resolved holoenzyme domains, with the hub domain represented as blue outline and kinase domains as yellow circles. (C and D) *Top*, representative EM images showing apparent high-order clustering of holoenzymes (3 or more) for CaMKIIα and CaMKIIβ, respectively (dotted outlines). *Bottom*, annotated representation of resolved holoenzyme domains, as in panels (A) and (B), with holoenzyme clusters indicated by dotted outline. Scale bar = 50 nm in panels (A – D). (E) Percentage of paired/clustered holoenzymes as assigned by visual inspection of micrographs for CaMKIIα and CaMKIIβ image datasets (n=1,207 particles for CaMKIIα and 1,255 particles for CaMKIIβ), indicate an approximately equal propensity for holoenzyme clustering between these two isoforms under basal-state conditions.

**Figure S4.**
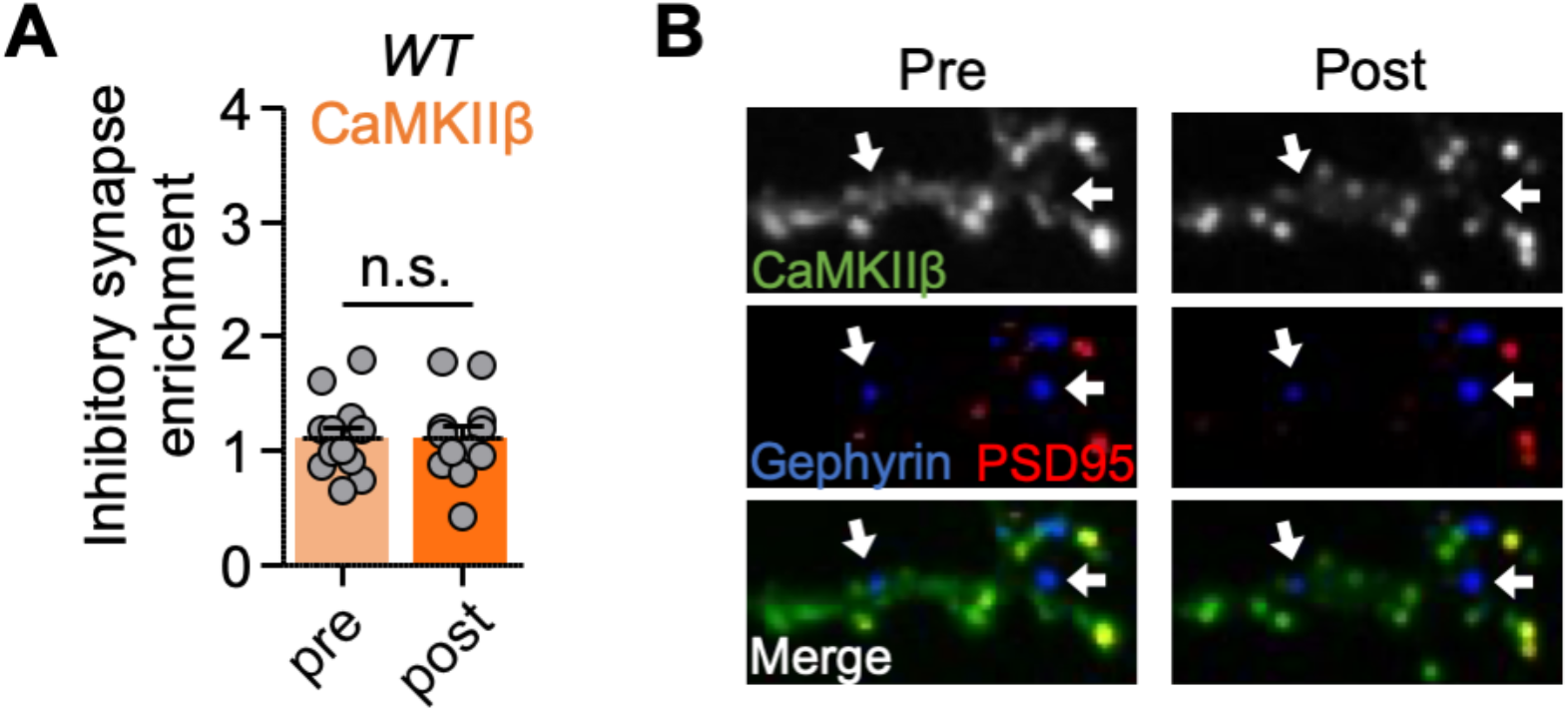
CaMKIIβ does not move to inhibitory synapses following prolonged glutamate. Quantifications show mean ± SEM. (A) Quantification of inhibitory synapse enrichment induced by excitotoxic glutamate (100 μM glutamate, 5 min) in WT cultured hippocampal neurons (paired two-tailed t-test: p=0.4689). (B) Representative confocal images show overexpressed CaMKIIβ, endogenous PSD95 (in red) to mark excitatory synapses, and endogenous gephyrin (in blue) to mark inhibitory synapses.

**Figure S5.**
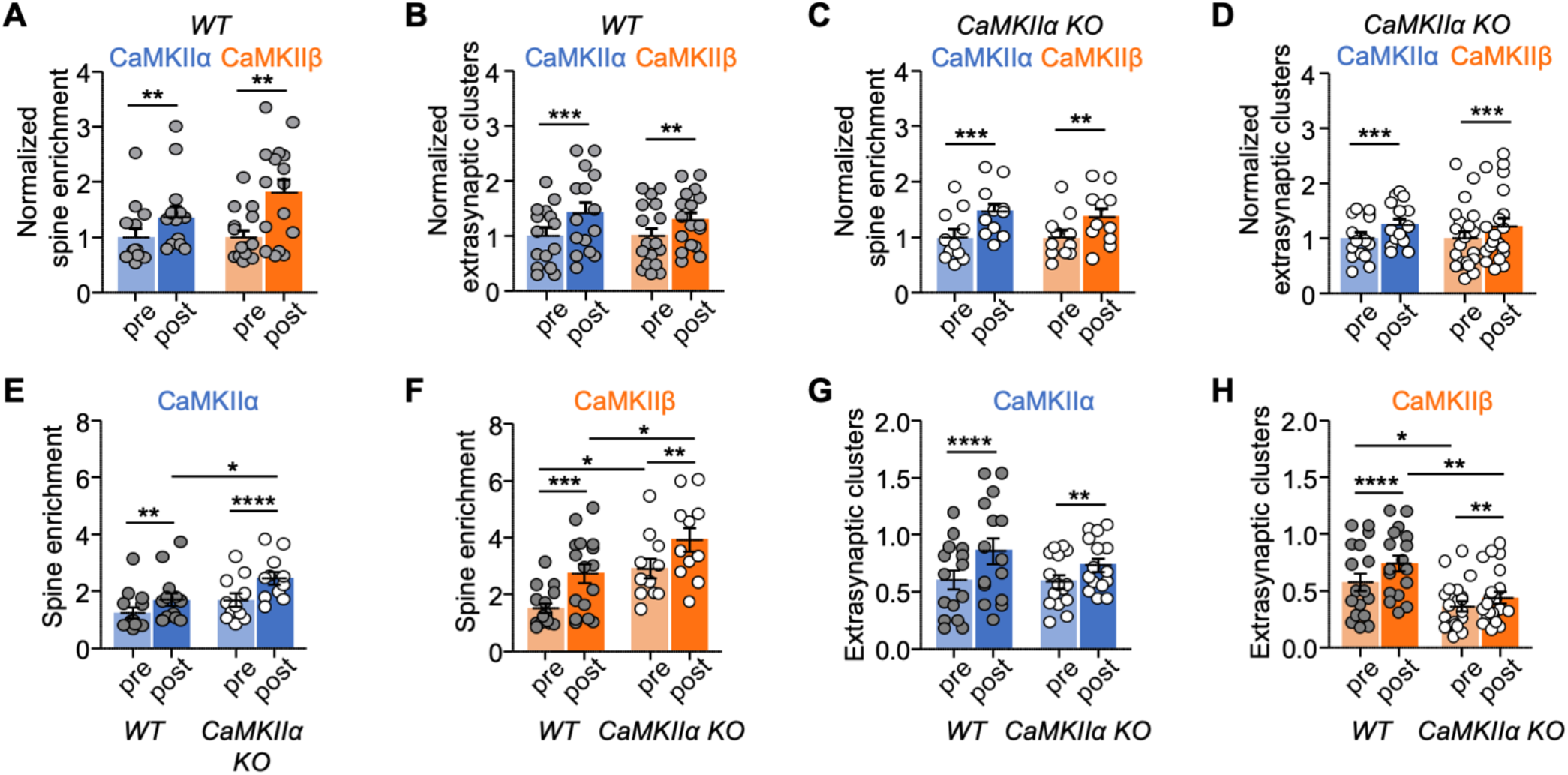
CaMKII movement normalized and compared by genotype. Panels A-D show CaMKIIα vs. CaMKIIβ isoform comparison after normalizing to baseline. Panels E-H show WT vs. CaMKIIα KO genotype comparison. Quantifications show mean ± SEM. *p<0.05, **p<0.01, ***p<0.001, ****p<0.0001. (A) CaMKIIα vs. CaMKIIβ movement to excitatory synapses in WT (two-way repeated-measures ANOVA, Bonferroni’s test: **p=0.0012 for pre vs. post in CaMKIIα; **p=0.0017 for pre vs. post in CaMKIIβ). (B) CaMKIIα vs. CaMKIIβ clustering at extra-synaptic cluster sites in WT (two-way repeated-measures ANOVA, Bonferroni’s test: ***p=0.0001 for pre vs. post in CaMKIIα; **p=0.0035 for pre vs. post in CaMKIIβ). (C) CaMKIIα vs. CaMKIIβ movement to excitatory synapses in CaMKIIα KO (two-way repeated-measures ANOVA, Bonferroni’s test: ***p=0.0001 for pre vs. post in CaMKIIα; **p=0.0017 for pre vs. post in CaMKIIβ). (D) CaMKIIα vs. CaMKIIβ clustering at extra-synaptic cluster sites in CaMKIIα KO (two-way repeated-measures ANOVA, Bonferroni’s test: **p=0.0004 for pre vs. post in CaMKIIα; **p=0.0007 for pre vs. post in CaMKIIβ). (E) CaMKIIα movement to excitatory synapses in WT vs. CaMKIIα KO (two-way repeated-measures ANOVA, Bonferroni’s test: **p=0.0018 for pre vs. post in WT; ****p<0.0001 for pre vs. post in CaMKIIα KO; *p=0.0344 for post WT vs. post CaMKIIα KO). (F) CaMKIIβ movement to excitatory synapses in WT vs. CaMKIIα KO (two-way repeated-measures ANOVA, Bonferroni’s test: ****p<0.0001 for pre vs. post in WT; **p=0.0024 for pre vs. post in CaMKIIα KO; *p=0.0230 for pre WT vs. pre CaMKIIα KO; *p=0.0078 for post WT vs. post CaMKIIα KO). (G) CaMKIIα extra-synaptic clustering in WT vs. CaMKIIα KO (two-way repeated-measures ANOVA, Bonferroni’s test: ****p<0.0001 for pre vs. post in WT; *p=0.0179 for pre vs. post in CaMKIIα KO). (H) CaMKIIβ extra-synaptic clustering in WT vs. CaMKIIα KO (two-way repeated-measures ANOVA, Bonferroni’s test: ****p<0.0001 for pre vs. post in WT; **p=0.0058 for pre vs. post in CaMKIIα KO; *p=0.0303 for pre WT vs. pre CaMKIIα KO; **p=0.0015 for post WT vs. post CaMKIIα KO).

## REFERENCES

Aronowski, J., Grotta, J.C., and Waxham, M.N. (1992). Ischemia-induced translocation of Ca2+/calmodulin-dependent protein kinase II: potential role in neuronal damage. J Neurochem 58, 1743–1753.

Banani, S.F., Lee, H.O., Hyman, A.A., and Rosen, M.K. (2017). Biomolecular condensates: organizers of cellular biochemistry. Nat Rev Mol Cell Biol 18, 285–298.

Barcomb, K., Hell, J.W., Benke, T.A., and Bayer, K.U. (2016). The CaMKII/GluN2B Protein Interaction Maintains Synaptic Strength. J Biol Chem 291, 16082–16089.

Barria, A., and Malinow, R. (2005). NMDA receptor subunit composition controls synaptic plasticity by regulating binding to CaMKII. Neuron 48, 289–301.

Bayer, K.U., De Koninck, P., Leonard, A.S., Hell, J.W., and Schulman, H. (2001). Interaction with the NMDA receptor locks CaMKII in an active conformation. Nature 411, 801–805.

Bayer, K.U., De Koninck, P., and Schulman, H. (2002). Alternative splicing modulates the frequency-dependent response of CaMKII to Ca(2+) oscillations. The EMBO journal 21, 3590–3597.

Bayer, K.U., LeBel, E., McDonald, G.L., O’Leary, H., Schulman, H., and De Koninck, P. (2006). Transition from reversible to persistent binding of CaMKII to postsynaptic sites and NR2B. J Neurosci 26, 1164–1174.

Bayer, K.U., Lohler, J., Schulman, H., and Harbers, K. (1999). Developmental expression of the CaM kinase II isoforms: ubiquitous gamma- and delta-CaM kinase II are the early isoforms and most abundant in the developing nervous system. Brain research Molecular brain research 70, 147–154.

Bayer, K.U., and Schulman, H. (2019). CaM Kinase: Still Inspiring at 40. Neuron 103, 380–394.

Bennett, M.K., and Kennedy, M.B. (1987). Deduced primary structure of the beta subunit of brain type II Ca2+/calmodulin-dependent protein kinase determined by molecular cloning. Proc Natl Acad Sci U S A 84, 1794–1798.

Bhattacharyya, M., Karandur, D., and Kuriyan, J. (2020a). Structural Insights into the Regulation of Ca(2+)/Calmodulin-Dependent Protein Kinase II (CaMKII). Cold Spring Harb Perspect Biol 12.

Bhattacharyya, M., Lee, Y.K., Muratcioglu, S., Qiu, B., Nyayapati, P., Schulman, H., Groves, J.T., and Kuriyan, J. (2020b). Flexible linkers in CaMKII control the balance between activating and inhibitory autophosphorylation. Elife 9.

Bhattacharyya, M., Stratton, M.M., Going, C.C., McSpadden, E.D., Huang, Y., Susa, A.C., Elleman, A., Cao, Y.M., Pappireddi, N., Burkhardt, P., et al. (2016). Molecular mechanism of activation-triggered subunit exchange in Ca(2+)/calmodulin-dependent protein kinase II. Elife 5.

Bradshaw, J.M., Hudmon, A., and Schulman, H. (2002). Chemical quenched flow kinetic studies indicate an intraholoenzyme autophosphorylation mechanism for Ca2+/calmodulin-dependent protein kinase II. J Biol Chem 277, 20991–20998.

Brangwynne, C.P., Eckmann, C.R., Courson, D.S., Rybarska, A., Hoege, C., Gharakhani, J., Julicher, F., and Hyman, A.A. (2009). Germline P granules are liquid droplets that localize by controlled dissolution/condensation. Science 324, 1729–1732.

Brocke, L., Chiang, L.W., Wagner, P.D., and Schulman, H. (1999). Functional implications of the subunit composition of neuronal CaM kinase II. J Biol Chem 274, 22713–22722.

Buonarati, O.R., Cook, S.G., Goodell, D.J., Chalmers, N., Rumian, N.L., Tullis, J.E., Restrepo, S., Coultrap, S.J., Quillinan, N., Herson, P.S., et al. (2020). CaMKII versus DAPK1 binding to GluN2B in ischemic neuronal cell death after resuscitation from cardiac arrest. Cell Rep 30, 1–8.

Chao, L.H., Stratton, M.M., Lee, I.H., Rosenberg, O.S., Levitz, J., Mandell, D.J., Kortemme, T., Groves, J.T., Schulman, H., and Kuriyan, J. (2011). A mechanism for tunable autoinhibition in the structure of a human Ca2+/calmodulin-dependent kinase II holoenzyme. Cell 146, 732–745.

Chen, X., Wu, X., Wu, H., and Zhang, M. (2020). Phase separation at the synapse. Nat Neurosci 23, 301–310.

Colbran, R.J. (1993). Inactivation of Ca2+/calmodulin-dependent protein kinase II by basal autophosphorylation. J Biol Chem 268, 7163–7170.

Cook, S.G., Bourke, A.M., O’Leary, H., Zaegel, V., Lasda, E., Mize-Berge, J., Quillinan, N., Tucker, C.L., Coultrap, S.J., Herson, P.S., et al. (2018). Analysis of the CaMKIIalpha and beta splice-variant distribution among brain regions reveals isoform-specific differences in holoenzyme formation. Sci Rep 8, 5448.

Cook, S.G., Goodell, D.J., Restrepo, S., Arnold, D.B., and Bayer, K.U. (2019). Simultaneous live-imaging of multiple endogenous proteins reveals a mechanism for Alzheimer’s-related plasticity impairment. Cell Rep 27, 658–665.

Coultrap, S.J., and Bayer, K.U. (2011). Improving a natural CaMKII inhibitor by random and rational design. PLoS One 6, e25245.

Coultrap, S.J., and Bayer, K.U. (2012a). Ca^2+^/Calmodulin-Dependent Protein Kinase II (CaMKII). In Neuromethods: Protein Kinase Technologies, H. Mukai, ed. (Springer), pp. 49–72.

Coultrap, S.J., and Bayer, K.U. (2012b). CaMKII regulation in information processing and storage. Trends in neurosciences 35, 607–618.

Coultrap, S.J., and Bayer, K.U. (2014). Nitric oxide induces Ca2+-independent activity of the Ca2+/calmodulin-dependent protein kinase II (CaMKII). J Biol Chem 289, 19458–19465.

Coultrap, S.J., Buard, I., Kulbe, J.R., Dell’Acqua, M.L., and Bayer, K.U. (2010). CaMKII autonomy is substrate-dependent and further stimulated by Ca2+/calmodulin. J Biol Chem 285, 17930–17937.

Coultrap, S.J., Freund, R.K., O’Leary, H., Sanderson, J.L., Roche, K.W., Dell’Acqua, M.L., and Bayer, K.U. (2014). Autonomous CaMKII mediates both LTP and LTD using a mechanism for differential substrate site selection. Cell Reports 6, 431–437.

Coultrap, S.J., Vest, R.S., Ashpole, N.M., Hudmon, A., and Bayer, K.U. (2011). CaMKII in cerebral ischemia. Acta Pharmacol Sin 32, 861–872.

De Koninck, P., and Schulman, H. (1998). Sensitivity of CaM kinase II to the frequency of Ca2+ oscillations. Science 279, 227–230.

Deng, G., Orfila, J.E., Dietz, R.M., Moreno-Garcia, M., Rodgers, K.M., Coultrap, S.J., Quillinan, N., Traystman, R.J., Bayer, K.U., and Herson, P.S. (2017). Autonomous CaMKII Activity as a Drug Target for Histological and Functional Neuroprotection after Resuscitation from Cardiac Arrest. Cell Rep 18, 1109–1117.

Dosemeci, A., Reese, T.S., Petersen, J., and Tao-Cheng, J.H. (2000). A novel particulate form of Ca(2+)/calmodulin-dependent [correction of Ca(2+)/CaMKII-dependent] protein kinase II in neurons. J Neurosci 20, 3076–3084.

Dyla, M., and Kjaergaard, M. (2020). Intrinsically disordered linkers control tethered kinases via effective concentration. Proc Natl Acad Sci U S A 117, 21413–21419.

Flory, P.J. (1975). Spatial configuration of macromolecular chains. Science 188, 1268–1276.

Giese, K.P., Fedorov, N.B., Filipkowski, R.K., and Silva, A.J. (1998). Autophosphorylation at Thr286 of the alpha calcium-calmodulin kinase II in LTP and learning. Science 279, 870–873.

Gross, G.G., Junge, J.A., Mora, R.J., Kwon, H.B., Olson, C.A., Takahashi, T.T., Liman, E.R., Ellis-Davies, G.C., McGee, A.W., Sabatini, B.L., et al. (2013). Recombinant probes for visualizing endogenous synaptic proteins in living neurons. Neuron 78, 971–985.

Halt, A.R., Dallpiazza, R.F., Zhou, Y., Stein, I.S., Qian, H., Juntti, S., Wojcik, S., Brose, N., Silva, A.J., and Hell, J.W. (2012). CaMKII binding to GluN2B is critical during memory consolidation. EMBO J 31, 1203–1216.

Hanson, P.I., Meyer, T., Stryer, L., and Schulman, H. (1994). Dual role of calmodulin in autophosphorylation of multifunctional CaM kinase may underlie decoding of calcium signals. Neuron 12, 943–956.

Hell, J.W. (2014). CaMKII: claiming center stage in postsynaptic function and organization. Neuron 81, 249–265.

Hoelz, A., Nairn, A.C., and Kuriyan, J. (2003). Crystal structure of a tetradecameric assembly of the association domain of Ca2+/calmodulin-dependent kinase II. Mol Cell 11, 1241–1251.

Hosokawa, T., Liu, P.-W., Cai, Q., Ferreira, J.S., Levet, F., Butler, C., Sibarita, J.-B., Choquet, D., Groc, L., Hosy, E., et al. (2020). CaMKII activation triggers persistent formation and segregation of postsynaptic liquid phase. bioRxiv, 2020.2011.2025.397091.

Hudmon, A., Aronowski, J., Kolb, S.J., and Waxham, M.N. (1996). Inactivation and self-association of Ca2+/calmodulin-dependent protein kinase II during autophosphorylation. J Biol Chem 271, 8800–8808.

Hudmon, A., Kim, S.A., Kolb, S.J., Stoops, J.K., and Waxham, M.N. (2001). Light scattering and transmission electron microscopy studies reveal a mechanism for calcium/calmodulin-dependent protein kinase II self-association. J Neurochem 76, 1364–1375.

Hudmon, A., Lebel, E., Roy, H., Sik, A., Schulman, H., Waxham, M.N., and De Koninck, P. (2005). A mechanism for Ca2+/calmodulin-dependent protein kinase II clustering at synaptic and nonsynaptic sites based on self-association. J Neurosci 25, 6971–6983.

Incontro, S., Diaz-Alonso, J., Iafrati, J., Vieira, M., Asensio, C.S., Sohal, V.S., Roche, K.W., Bender, K.J., and Nicoll, R.A. (2018). The CaMKII/NMDA receptor complex controls hippocampal synaptic transmission by kinase-dependent and independent mechanisms. Nat Commun 9, 2069.

Kanaseki, T., Ikeuchi, Y., Sugiura, H., and Yamauchi, T. (1991). Structural features of Ca2+/calmodulin-dependent protein kinase II revealed by electron microscopy. J Cell Biol 115, 1049–1060.

Karandur, D., Bhattacharyya, M., Xia, Z., Lee, Y.K., Muratcioglu, S., McAffee, D., McSpadden, E. D., Qiu, B., Groves, J.T., Williams, E.R., et al. (2020). Breakage of the oligomeric CaMKII hub by the regulatory segment of the kinase. Elife 9.

Li, P., Banjade, S., Cheng, H.C., Kim, S., Chen, B., Guo, L., Llaguno, M., Hollingsworth, J.V., King, D.S., Banani, S.F., et al. (2012). Phase transitions in the assembly of multivalent signalling proteins. Nature 483, 336–340.

Lisman, J., Yasuda, R., and Raghavachari, S. (2012). Mechanisms of CaMKII action in longterm potentiation. Nature reviews Neuroscience 13, 169–182.

McSpadden, E.D., Xia, Z., Chi, C.C., Susa, A.C., Shah, N.H., Gee, C.L., Williams, E.R., and Kuriyan, J. (2019). Variation in assembly stoichiometry in non-metazoan homologs of the hub domain of Ca(2+) /calmodulin-dependent protein kinase II. Protein Sci 28, 1071–1082.

Miller, S.G., and Kennedy, M.B. (1986). Regulation of brain type II Ca2+/calmodulin-dependent protein kinase by autophosphorylation: a Ca2+-triggered molecular switch. Cell 44, 861–870.

Mora, R.J., Roberts, R.W., and Arnold, D.B. (2013). Recombinant probes reveal dynamic localization of CaMKIIalpha within somata of cortical neurons. J Neurosci 33, 14579–14590.

Myers, J.B., Zaegel, V., Coultrap, S.J., Miller, A.P., Bayer, K.U., and Reichow, S.L. (2017). The CaMKII holoenzyme structure in activation-competent conformations. Nat Commun 8, 15742.

Nguyen, T.A., Sarkar, P., Veetil, J.V., Davis, K.A., Puhl, H.L., 3rd, and Vogel, S.S. (2015). Covert Changes in CaMKII Holoenzyme Structure Identified for Activation and Subsequent Interactions. Biophys J 108, 2158–2170.

Nguyen, T.A., Sarkar, P., Veetil, J.V., Koushik, S.V., and Vogel, S.S. (2012). Fluorescence polarization and fluctuation analysis monitors subunit proximity, stoichiometry, and protein complex hydrodynamics. PLoS One 7, e38209.

Rellos, P., Pike, A.C., Niesen, F.H., Salah, E., Lee, W.H., von Delft, F., and Knapp, S. (2010). Structure of the CaMKIIdelta/calmodulin complex reveals the molecular mechanism of CaMKII kinase activation. PLoS Biol 8, e1000426.

Rich, R.C., and Schulman, H. (1998). Substrate-directed function of calmodulin in autophosphorylation of Ca2+/calmodulin-dependent protein kinase II. J Biol Chem 273, 28424–28429.

Rosenberg, O.S., Deindl, S., Sung, R.J., Nairn, A.C., and Kuriyan, J. (2005). Structure of the autoinhibited kinase domain of CaMKII and SAXS analysis of the holoenzyme. Cell 123, 849–860.

Rossetti, T., Banerjee, S., Kim, C., Leubner, M., Lamar, C., Gupta, P., Lee, B., Neve, R., and Lisman, J. (2017). Memory Erasure Experiments Indicate a Critical Role of CaMKII in Memory Storage. Neuron 96, 207–216 e202.

Sanhueza, M., Fernandez-Villalobos, G., Stein, I.S., Kasumova, G., Zhang, P., Bayer, K.U., Otmakhov, N., Hell, J.W., and Lisman, J. (2011). Role of the CaMKII/NMDA receptor complex in the maintenance of synaptic strength. J Neurosci 31, 9170–9178.

Scheres, S.H. (2012). RELION: implementation of a Bayesian approach to cryo-EM structure determination. J Struct Biol 180, 519–530.

Shen, K., and Meyer, T. (1999). Dynamic control of CaMKII translocation and localization in hippocampal neurons by NMDA receptor stimulation. Science 284, 162–166.

Singla, S.I., Hudmon, A., Goldberg, J.M., Smith, J.L., and Schulman, H. (2001). Molecular characterization of calmodulin trapping by calcium/calmodulin-dependent protein kinase II. J Biol Chem 276, 29353–29360.

Sloutsky, R., Dziedzic, N., Dunn, M.J., Bates, R.M., Torres-Ocampo, A.P., Boopathy, S., Page, B., Weeks, J.G., Chao, L.H., and Stratton, M.M. (2020). Heterogeneity in human hippocampal CaMKII transcripts reveals allosteric hub-dependent regulation. Sci Signal 13.

Strack, S., McNeill, R.B., and Colbran, R.J. (2000). Mechanism and regulation of calcium/calmodulin-dependent protein kinase II targeting to the NR2B subunit of the N-methyl-D-aspartate receptor. J Biol Chem 275, 23798–23806.

Stratton, M., Lee, I.H., Bhattacharyya, M., Christensen, S.M., Chao, L.H., Schulman, H., Groves, J.T., and Kuriyan, J. (2014). Activation-triggered subunit exchange between CaMKII holoenzymes facilitates the spread of kinase activity. Elife 3, e01610.

Tang, G., Peng, L., Baldwin, P.R., Mann, D.S., Jiang, W., Rees, I., and Ludtke, S.J. (2007). EMAN2: an extensible image processing suite for electron microscopy. J Struct Biol 157, 38–46.

Thaler, C., Koushik, S.V., Puhl, H.L., 3rd, Blank, P.S., and Vogel, S.S. (2009). Structural rearrangement of CaMKIIalpha catalytic domains encodes activation. Proc Natl Acad Sci U S A 106, 6369–6374.

Tombes, R.M., Faison, M.O., and Turbeville, J.M. (2003). Organization and evolution of multifunctional Ca(2+)/CaM-dependent protein kinase genes. Gene 322, 17–31.

Vest, R.S., O’Leary, H., and Bayer, K.U. (2009). Differential regulation by ATP versus ADP further links CaMKII aggregation to ischemic conditions. FEBS Lett 583, 3577–3581.

Vest, R.S., O’Leary, H., Coultrap, S.J., Kindy, M.S., and Bayer, K.U. (2010). Effective post-insult neuroprotection by a novel Ca(2+)/ calmodulin-dependent protein kinase II (CaMKII) inhibitor. J Biol Chem 285, 20675–20682.

Woodgett, J.R., Davison, M.T., and Cohen, P. (1983). The calmodulin-dependent glycogen synthase kinase from rabbit skeletal muscle. Purification, subunit structure and substrate specificity. Eur J Biochem 136, 481–487.

Zivanov, J., Nakane, T., Forsberg, B.O., Kimanius, D., Hagen, W.J., Lindahl, E., and Scheres, S.H. (2018). New tools for automated high-resolution cryo-EM structure determination in RELION-3. Elife 7.

